# Surviving on limited resources: effects of caloric restriction on growth, gene expression and gut microbiota in a species with male pregnancy (*Hippocampus erectus*)

**DOI:** 10.1101/2023.10.05.560864

**Authors:** Freya Adele Pappert, Vincent Alexander Wüst, Carmen Fontanes Eguiguren, Olivia Roth

**Affiliations:** Marine Evolutionary Biology, Zoological Institute, Christian-Albrechts-Universität Kiel, Am Botanischen Garten 1-9, 24118 Kiel, Germany; Evolutionary Ecology of Marine Fishes, Helmholtz-Centre for Ocean Research Kiel (GEOMAR), 24105 Kiel, Germany; University of Vienna, Djerassipl. 1, 1030 Vienna, Austria

**Keywords:** caloric restriction, resource allocation, life-history, seahorse, male pregnancy

## Abstract

Caloric restriction (CR) studies have traditionally focused on species with conventional reproductive roles, emphasizing female’s greater investment in costly gametes and parental care. While the divergent impact of CR on males and females is evident across species, the factors driving this variation, i.e., resource allocation to reproductive elements as part of distinct life history strategies, remain unclear. To address this, we investigated the effects of CR on development, gene expression, and intestinal microbiota in the lined seahorse *Hippocampus erectus,* a species with male pregnancy, where fathers invest in offspring through gestation. Juvenile seahorses were subjected to ad libitum (AL) or CR feeding for 5 months. CR stunted male growth and brood pouch development, reflecting the energy demands of this crucial parental care trait. However, condition index declined in CR females but not males, while ovarian weight remained unchanged. Transcriptome analysis demonstrated organ- and sex-specific responses to CR with distinct lipid and energy-related pathways activated in male and female livers, indicative of survival enhancement strategies. CR had minimal impact on genes associated with spermatogenesis, but downregulated lipid metabolic and inflammatory genes in ovaries, emphasizing the importance of pre-copulatory resource allocation in female gametes. CR strongly shaped gut microbial composition, creating distinct communities from AL seahorses while also driving sex-specific taxonomic differences. Our research indicates that nutrient limitatiońs impact on males and females is influenced by their allocation of resources to reproduction and parental investment. We underscore the significance of studying species with diverse reproductive strategies, sex roles, and life-history strategies.

## 1. Introduction

Females and males have distinct reproductive strategies, leading to specific energy allocation. Females typically invest more in reproduction than males, often through a combination of costlier games and greater parental care (1). For example, in many taxa, females produce energetically expensive eggs and bear additional burdens, such as pregnancy and lactation (2,3). As direct reproductive investment is often greater in females due to the combined costs of gamete production and parental care, males in many species allocate more energy to direct activities that enhance reproductive success, such as courting and mate guarding (4,5). These differences in reproductive strategies likely influence how males and females allocate resources and energy in response to environmental challenges, such as caloric restriction (CR).

Studies have shown that female mice and fruit flies tend to react stronger to CR in terms of improved health- and lifespan compared to their male counterparts (6,7). In contrast, fasting extended lifespan of male but not female killifish, potentially facilitated by sex-specific expression changes (8). Despite these species-specific sex differences, biomedical research has historically neglected sex differences in CR responses, underlining that the drivers of this sexual dimorphism are still poorly understood (9,10).

CR induces a state of hormetic stress, stimulating a scarcity of food and prompting it to conserve energy until more nutrients become available (11). According to Kirkwood’s disposable soma theory, longevity requires investing in somatic maintenance, which takes priority over reproduction when resources are scarce (12). By postponing reproduction during periods of limited energy availability, animals can extend their lifespan within the constraints of their maximum biological potential, resuming breeding when resources become abundant again (13). This strategy aims to maximize overall reproductive success (14). Some species naturally have long periods of fasting during hibernations when food is scarce, including many reptiles and fish (15,16). However, species with long lifespans, high caloric diets and high reproductive outputs, such as queen bees, ants and termites challenge this theory (17,18). If done excessively, CR can have negative health impact in humans (e.g. extreme weight loss, lethargy, diminished libido etc.) (14,19–21).

Despite the general consensus that different species showcase varying sexual dimorphism with respect to the effects of CR, most research to date has focused predominantly on species with conventional reproductive roles, with female showing increased investment into both gametes and offspring (22). However, sex roles in the animal kingdom are not constrained and have evolved with flexibility across species (5,23,24), leaving a gap in our understanding regarding whether the sex-specific effects of fasting are primarily driven by differences in resource allocation to gamete production (e.g., egg vs. sperm) or by broader sexual reproductive roles, such as parental care. Understanding the ultimate drivers of sexual dimorphism in the effects of CR on health across different species holds significant implications for integrating concepts of sexual selection into medical sciences. This knowledge can offer valuable insights for CR interventions in sex- and gender-specific health or the development of CR mimetics, paving the way for improved strategies to enhance well-being and longevity (25).

To bridge the existing research gap, we designed an experiment focusing on how CR, implemented through intermittent fasting (IF), affects development, gut microbiota and aging pathways in a species with male pregnancy, the lined seahorse *Hippocampus erectus*. In this species the female transfers her eggs to the male’s pouch, which he then fertilizes with his sperm. The embryos are embedded in the fully enclosed pouch containing blood vessels supplying nutrients and oxygen, on top of providing a controlled environment (26,27). The unique reproductive biology of this species allows to disentangle the cost of investment into the brooding structure (male) vs. investment into egg production (female), which are two energetically costly traits typically found in females. This provides an opportunity to uncover how divergent life histories and reproductive trade-offs affect sexual dimorphic responses to CR.

CR influences signaling pathways involved in growth, metabolism, oxidative stress response, damage repair, inflammation, autophagy, and proteostasis, shaping the aging process and yielding positive physiological outcomes. The latter encompass improved cardiovascular health, reduced oxidative damage, enhanced DNA repair, promotion of autophagy, and mitigating gut dysbiosis (21,28,29). Lifelong CR increases lifespan in many species, including yeast (30), nematodes (31), fruit flies (32), and rodents (21,33). Rejuvenating properties were suggested for mice, rhesus monkey and humans fed on calorie-restricted diets who showed DNA methylation patterns equal to their younger counterparts (34), and a younger healthier microbial composition after two months of CR (35).

We took 52 three-month-old juvenile (not sexually mature) seahorses, which were divided equally into two feeding groups: Ad libitum (AL) and calorie-restricted (CR). Over the course of 5 months, the AL group was fed twice a day, while the CR group was fed every-other-day (also twice), resulting in a calorie restriction of approximately 50%. As seahorses lack a stomach, they are limited in the consumption of food at each food provisioning (36). This avoids overeating in the AL and ensures a restriction of calories in the fasting group. We measured the weight of the seahorses at four time points over the experimental period of 5 months. In males we examined physiological developmental differences in pouch formation, the male critical reproductive structure. In females we compared, at the end of fasting period, the ovary masses to assess egg production. Using RNA sequencing we analyzed transcriptome-wide differential gene expression from liver, head kidney, and gonads, between male vs. female AL and CR groups. We sampled hind-gut to genotype microbial composition using 16S rRNA amplicon sequencing to investigate potential sex-specific changes induced by fasting.

We further compared individuals exposed to CR to young and old control individuals (respectively 4 months and 3 years old) from the same stock populations, to better understand the effect of CR on senescence and provide insights into how CR-induced genetic and microbial modifications may influence aging and rejuvenation processes.

## 2. Material and methods

### 2.1 Study design and organism

Three-month-old juvenile *Hippocampus erectus* from an aquarium bred population kept at the GEOMAR Helmholtz Centre for Ocean Research Kiel were used for the experiment. Fifty-two juvenile seahorses were distributed across 18 tanks (20 l aquaria), with 16 tanks housing three seahorses each, and 2 tanks (one for the AL treatment and one for CR treatment) housing two seahorses each.

The aquaria had a recirculating system with a temperature of 23-25°C, salinity 30-35 PSU, nitrate 20-50 mg/l, phosphate below 1 ppm, ozonisation, and a 20 % weekly water change. Filters, plastic seagrass, and oxygen tubes were cleaned separately with hot water bi-weekly, left-over food or waste was suctioned and removed thirty minutes after feeding to prevent the biofilm formation.

We allocated twenty-six individuals to an AL feeding regime and another twenty-six to a CR regime. The tanks were divided into 9 AL and 9 CR, with alternating tanks placed next to each other to reduce tank location effects. All fish were fed with defrosted mysids, the AL group was fed twice a day, in the morning (8am) and in the late afternoon (4pm); the CR group was fed every other day, i.e., one day no food, the next day they were fed like the AL group. Seahorses show a secondary loss of stomach, requiring them to feed continuously for survival. They usually consume small crustaceans floating in the water, using their camouflage to remain still and ambush prey at the right moment with a swift strike (36).

To ensure that observed effects in the CR group were due to calorie reduction and not nutrient deficiency, both groups were given vitamins. The supplements were provided equally on the days when both groups were fed and included 15 µl of Microbe-Lift Garlic feed supplement (contains pure garlic, known to reduce bacterial diseases in fish (37)), 15 µl of JBL Atvitol multivitamin drops for aquarium fish and half a capsule (730mg) of Omega-3 1000 (unsaturated omega-3 fatty acids and vitamin E). These additives were left in the defrosted food for 2-3 minutes to ensure absorption. Seahorses were monitored daily during feeding for a health check and/or any abnormal swimming behaviours.

At the start of the experiment, all animals were weighed and measured for length before being placed in their respective tanks. For the first seven days, we gradually reduced the food in the CR group by feeding them only once in the morning on fasting days to minimize stress and facilitate habituation. Starting from day eight, the exact feeding regime was implemented, with the CR group receiving no mysids on alternate days. The study spanned a total of 157 days, which roughly equates to five months, during which, pouch development was closely monitored to determine when sexual maturation was taking place, and to ensure separation of males and females in same sex-tanks to prevent mating. For growth monitoring they were weighed in a small beaker, with previously calibrated tank water, roughly every 50 days. Upon termination of the dietary experiment, the seahorses were euthanized with an overdose of MS-222 (Tricaine methane sulfonate; 500 mg/l; Sigma-Aldrich). They were weighed and measured for total body length (snout to tail). Additionally, female ovaries were weighed and the width and length of pouch were measured for males. Liver, head kidney, and gonads for gene expression analysis were immediately preserved in RNA later stored at 4°C for three days, followed by long-term storage at -20°C. Hind-gut for microbiota genotyping was placed in a sterile Eppendorf tube and stored at - 80°C.

To assess how CR influences aging compared to the AL treatment, we took four-month-old juveniles labelled as control young (CY), and three-year-old adults labelled as control old (CO) that had been kept in the same GEOMAR stocks. They were also euthanized with 500 mg/l of MS-222, then measured for weight and body length and dissected to retrieve organs, which were stored as described above.

### 2.2 RNA and DNA extraction, library preparation and sequencing

RNA extraction of liver, head kidney, and gonads was performed using the RNeasy Mini Kit from Qiagen (Venlo, Netherlands) according to the manufacturer’s instructions. The concentration of the extracted RNA was measured using a Peqlab NanoDrop ND-1000 spectral photometer (Erlangen, Germany), and the samples were subsequently stored at -80°C. 120 Samples of 40 randomly selected individuals (5 males and 5 females from AL, CR, CO, CY, multiplied by the three tissues) were sent to BGI Tech Solutions in Hong Kong for library preparation and mRNA sequencing. For library preparation the DNBSEQ Eukaryotic Strand-specific mRNA library was utilized, and the sequencing of stranded mRNA was carried out on the DNBseq platform. The sequencing parameters included 150bp reads and 25M clean paired-end reads per sample.

To extract the microbiota from the hind-gut, DNA extraction was performed using the DNeasy Blood & Tissue Kit (QIAGEN, Germany) following the manufacturer’s protocol including a pre-treatment for Gram-positive bacteria with ameliorations (38). Subsequently, a 16S PCR was conducted to verify the success of the extraction. The library preparation for a total of 57 individuals was performed by the Institute for Experimental Medicine (UKSH, Campus Kiel). This group included 37 seahorses from the dietary experiment with an additionally 10 young and old individuals (5 AL, 11 CR, 5 CY, and 5 CO females, as well as 10 AL, 11 CR, 5 CY, and 5 CO males). Amplicon sequencing of the V3-V4 hypervariable region (341f/806r) was conducted using the Illumina MiSeq platform (Illumina, USA) with 2x 300-bp paired-end read settings at the IKMB Kiel.

### 2.3 Data analysis

#### 2.3.1 Morphology

In our experiment we complied stringently to daily animal welfare screening processes and seahorses were euthanized when scores were insufficient, resulting in a total of 38 individuals who survived until the conclusion of the experiment (day 157). Among them, the survivors encompassed 16 individuals belonging to the AL group with 11 AL males and 5 AL females, while 22 individuals belonged to the CR group with 11 CR males, and 11 CR females. At the beginning of the experiment, we could not divide the seahorses by sex as they were not sexually mature, resulting in a smaller number of AL females compared to the other groups.

For the morphology analysis, we included only the 38 individuals which survived to the end of the experiment, with an additional separate analysis comparing the diet groups to the young and old age groups of seahorses (5 CY, 5 CO both sexes). All the statistical analyses were done in Rstudio (v.4.2.2) (39).

We tested various morphological parameters for both dietary treatment (AL, CR) and age control group (CO, CY) on male and female, including weight (g) and total body length (cm), also Fulton’s condition factor (weight/length³) to better assess overall fish condition (40). Additionally, we calculated relative male pouch length (pouch length / total body length * 100) (Figs. 1D and S1H) and relative ovary weight (ovary weight / total body weight * 100) (Figs. 1C and S1G). Normality was assessed using a Shapiro-Wilk test. Depending on the results, we applied either a Kruskal-Wallis test (KW-test), followed by a Dunn’s post hoc test, or a two-way ANOVA followed by a Tukey Honest Significant Difference (Tukey HSD) test ((Treatment × Sex), Table S6 for statistics). Furthermore, since weight measurements were taken at four time points (Days 1, 55, 111, and 157), we analysed growth trajectories over time using a linear mixed model, with individuals as a random effect: lmer(weight (g) ∼ Time × Treatment × Sex + (1|Ind), data = data). Pairwise comparisons for Treatment × Sex at each time point were conducted using *emmeans* (Table S6, Fig. 1A).

**Figure 1:**
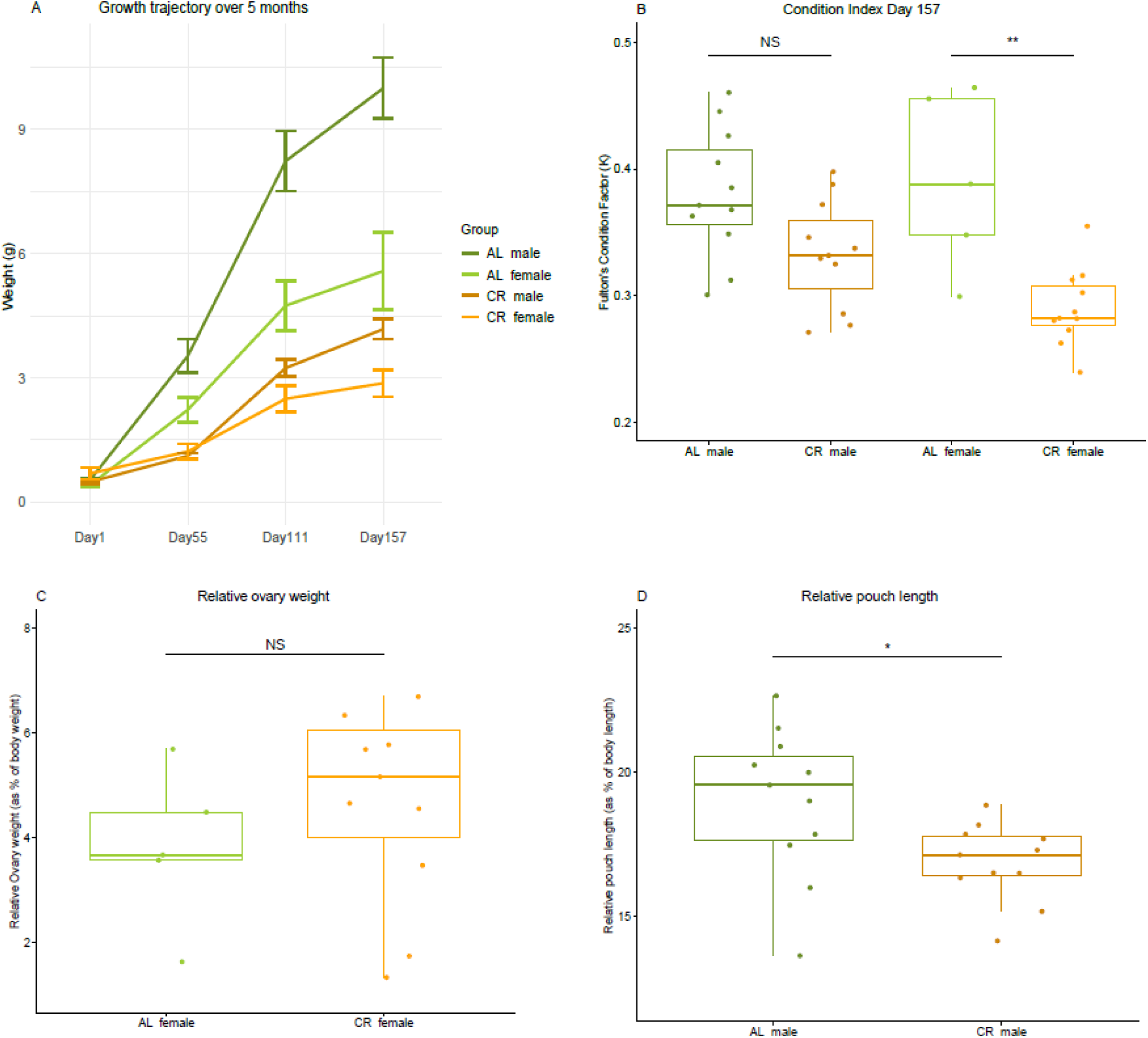
Effects of caloric restriction (CR) and ad libitum (AL) diets on growth, condition, and reproductive morphology in male and female seahorses. (A) Growth trajectories of male and female seahorses under CR and AL diets over five experimental months. The y-axis represents weight (grams), and the x-axis represents time with four measurement timepoints. AL males (dark green), AL females (light green), CR males (brown), and CR females (orange) are shown, with error bars indicating standard errors. (B) Condition index at the end of the experiment, comparing Fulton’s condition factor (weight/length³) across sexes and dietary treatments. Significant differences are denoted as *p < 0.05, **p < 0.001, and ***p < 0.0001. (C) Relative ovarian weight of CR and AL females, expressed as a percentage of body weight. The y-axis shows relative ovarian weight (%). (D) Relative pouch length in male seahorses from CR and AL treatments, expressed as a percentage of body length. The y-axis shows relative pouch length (%).

#### 2.3.2 Differential gene expression analysis

The fasting effects were characterized by constructing differential expression signatures associated with each organ. The integrity and quality of the RNA Illumina sequencing results from BGI Tech Solution were thoroughly assessed and controlled using FastQC (v.0.11.9) (41). The adapters had already been trimmed by BGI Tech Solutions. Regrettably, during the run, six samples failed, all of which were ovarian tissue (two AL, one CR, two OC, and one YC). For the alignment process, we used STAR (v.2.7.9) (42) to map the reads to a genome assembly of *Hippocampus erectus* (BioProject: PRJNA347499). Subsequently, read counts were obtained using TPMCalculator (43). In order to investigate potential distinctions among treatment groups (AL, CR, CO, CY) within specific organs, we partitioned the dataset by organ type. On logarithmically transformed count data we employed the “adonis2” function and the “bray” method, and conducted a PERMANOVA analysis (Permutational Multivariate Analysis of Variance Using Distance Matrices). Our examination evaluated the influences of diet and sex in the liver, head kidney, ovaries and testes. We did a principal component analysis (PCA) with multiple PCs to visualise group clustering and possible outliers (Figs. 2 and S2). For differential gene expression (DGE) analysis, we first used the edgeR package (v.3.40.2) (44), to scale the count data to counts per million (cpm) and filtered it based on a minimum threshold of 10 counts in at least 5 libraries.

**Figure 2:**
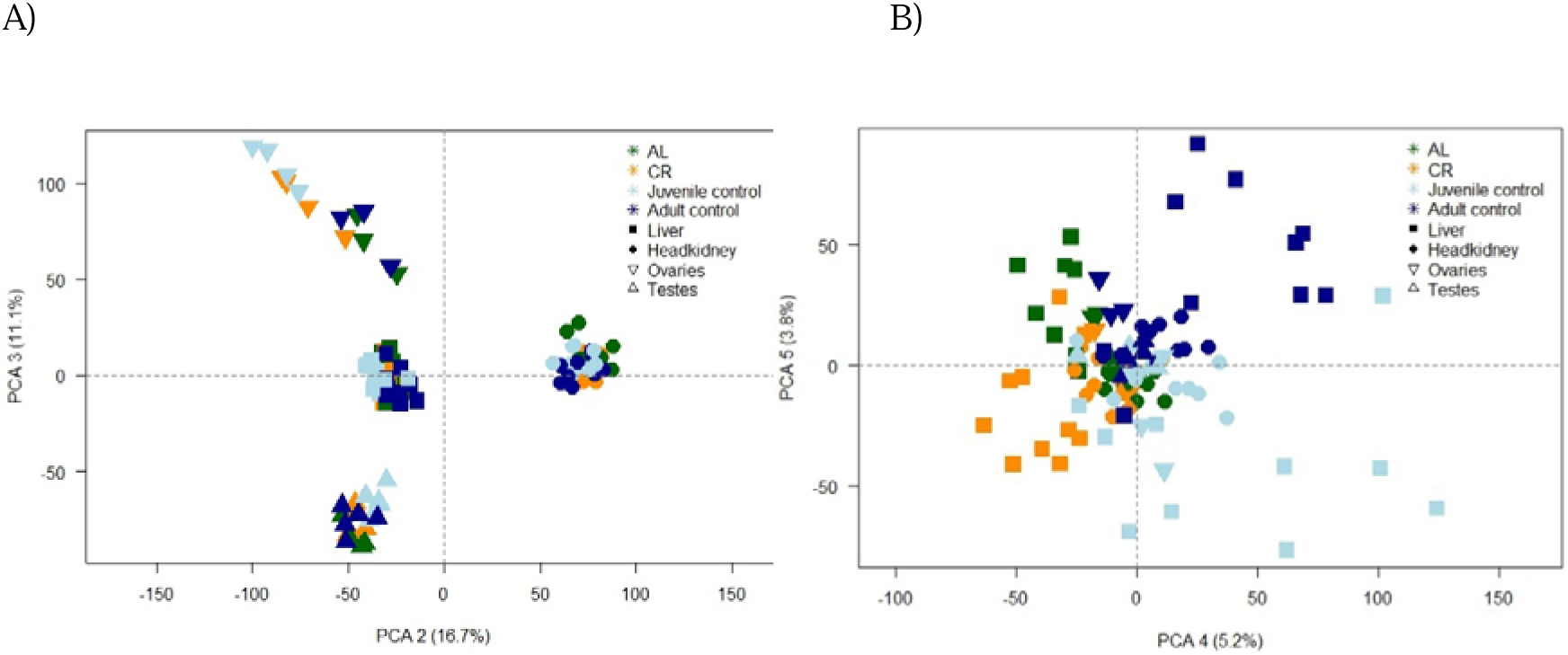
Principal Component Analysis (PCA) plots depicting organ separation and clustering of treatment groups. A) PCA plot of PC2 (16.7% variance) and PC3 (11.1% variance). PC2 separates the head kidney from the liver and gonads, while PC3 further differentiates male and female gonads. Groups are color-coded as follows: CR (orange), AL (green), juvenile control (light blue), and adult control (dark blue). Organs are represented by different symbols: liver (square), head kidney (circle), ovaries (downward triangle), and testes (upward triangle). (B) PCA plot of PC4 (5.2% variance) and PC5 (3.8% variance). PC4 separates treatment groups and age cohorts, while PC5 distinguishes CR and CY from AL and CO.

We performed this process separately for each organ, resulting in different gene sets for each: 13,015 genes for the liver, 16,470 genes for the head kidney, 17,411 genes for the testes, and 16,897 genes for the ovaries. To address composition biases, we normalized the data using the trimmed mean of values (TMM) with the calcNormFactors function. This step calculated normalization factors for each sample, effectively eliminating composition biases between libraries. Next, we utilized the limma package (v.3.54.1) (45) for DGE analysis. Limma is a linear model-based method that employs an empirical Bayes approach and the voom method to convert count data to a continuous scale accounting for batch effects and reducing technical variation. The voom method is well-suited for handling library sizes that vary significantly (46). To compare several groups, we created a matrix of independent contrasts, enabling us to perform a one-way analysis of deviance (ANODEV) for each gene. Subsequently, we estimated coefficients of interest between groups to determine log fold changes using the “contrasts.fit” function. We applied empirical Bayes moderation to shrink the estimated variance and performed a moderated t-test to identify differentially expressed genes (DEGs). For further downstream analysis, we focused only on adjusted P-values below 0.05 and corrected for multiple testing with Benjamini-Hochberg (BH) correction.

#### 2.3.3 Gene Ontology analysis

Gene ontology (GO) analysis was performed using the g:Profiler web tool (accessed on July 6, 2023). This tool enabled us to conduct gene enrichment analysis by comparing the significantly DEGs between AL and CR seahorses across the different organs. To annotate the genes of *Hippocampus erectus*, we employed OrthoFinder and conducted a homology-based search using *Danio rerio* (GRCz11) as a reference species. Adjusted statistical significance was set for “g:SCS threshold” value at 0.05, focusing exclusively on annotated genes. We treated numeric IDs as “ENTREZGENE_ACC” and limited the data sources to GO biological processes (BP), as well as biological pathways from the Kyoto Encyclopaedia of Genes and Genomes (KEGG) and Reactome (REAC) databases. To enhance the specificity of our analysis, we applied a maximum term size of 1000 to exclude overly broad categories and prioritize more informative results.

#### 2.3.4 Microbiome data analysis

The 16S rRNA amplicon sequencing analysis was performed via QIIME2 platform (v.2022.8.3) (47). With dada2 the raw paired-end Illumina reads were demultiplexed and amplicon primers for 16S V3-V4 region were trimmed, quality was controlled by “denoising” sequences (in order to better discriminate between true sequence diversity and sequencing errors). Low-quality reads were filtered (manual trimming at forward read at 250 bp and reverse at 210 bp), forward and reverse samples were merged and possible chimeras removed. A phylogenetic tree was constructed using “Fasttree” (v.2.1.11) that infers approximately-maximum-likelihood based on the longest root (48). Interactive α rarefaction curves were computed for max_depth of 20’000 to determine whether the samples were sequenced deeply enough to capture all the bacterial community members. Samples were classified for the V3/V4 hypervariable region based on taxonomic groups using the SILVA v138 database and a naïve Bayes based classifier (49). The amplicon sequence variants (ASV) were filtered to remove sequences from chloroplasts and mitochondria and exported at genus level as an operational taxonomic unit (OTU) for further analysis in RStudio (v.2022.07.2). To reduce noise, we applied a filtering threshold, including only OTUs that were present in at least three samples across the entire dataset. The remaining samples consisted of 5 AL, 11 CR, 5 CY, and 5 CO females, and 10 AL, 11 CR, 4 CY, and 4 CO males. OTU counts were normalized across samples to ensure data comparability.

α-diversity was measured using the Shannon index to measure diversity within individual seahorses using the vegan package (v.2.6.4) (50). β-diversity was assessed using the Bray Curtis dissimilarity matrix (based on relative abundance) to measure differences between the individual group of seahorses (vegdist function). Hypothesis testing for both α-diversity was done with Kruskal-Wallis test followed by a post hoc Dunn test (Table S6), while β-diversity was performed using PERMANOVA with both diet and sex as factors (x ∼ diet*sex, data=data, permutations=999), preceded by testing for normality with a Shapiro-Wilk normality test. To compare groups, pairwise.adonis2 v0.4 (51) was used as a post-hoc test, with Benjamini-Hochberg (BH) correction for p-value adjustment to account for multiple testing. To visualize the results, we used boxplots to display α-diversity and non-metric multidimensional scaling (NMDS) for β-diversity. For the NMDS, we employed 95% confidence ellipses to represent group variability, with the choice of ellipses based on standard error due to the high variance observed in the β-diversity data. This approach allows for a clearer representation of group dispersion while accounting for the inherent variability in the data. We examined the OTUs using the *envfit* function from the vegan package (v.2.6.4) to fit the OTUs to the NMDS ordination space and calculate the correlation between the ordination scores of OTUs and the treatment variables. By employing permutation tests with 999 iterations, we obtained p-values to assess the significance of these correlations. We plotted the OTUs that significantly explained the NMDS Bray-Curtis ordination (p.val < 0.001).

Relative abundance was calculated for the phylum and genus taxonomic level (Figs. 4A and S4B). The genus *Vibrio* (family *Vibrionaceae*) exhibited the highest frequency and variation across samples (Fig. 4A, S4B). Given the dominance of *Vibrio*, we sought to examine the broader background microbial community by focusing on less abundant taxa. To achieve this, we conducted an additional analysis excluding *Vibrio*, renormalized the data, and applied a centered log-ratio (CLR) transformation to convert relative abundances into log-ratio values. This transformation mitigates the unit sum constraint, allowing for more accurate comparisons across samples. Following this pre-processing, all statistical tests were repeated (Table S6), and alpha and beta diversity metrics were recalculated.

**Figure 3:**
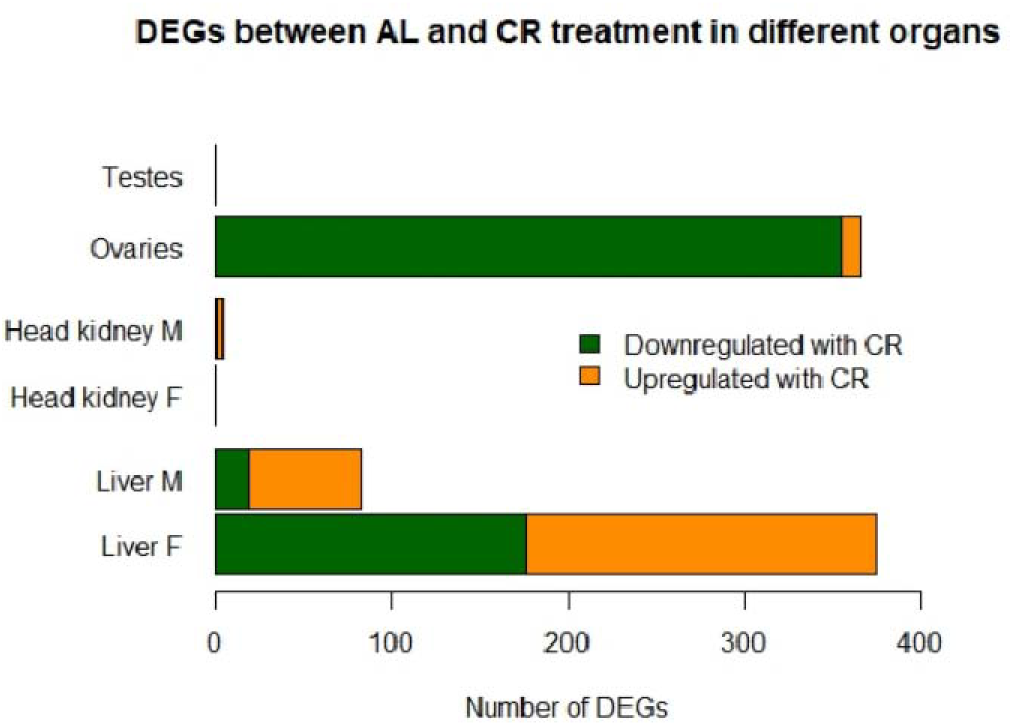
Sex-specific differential gene expression in AL and CR-fed seahorses across organs. Bar plot showing differentially expressed genes (DEGs) between AL and CR-fed seahorses across various organs, categorized by sex (M = male, F = female) on the y-axis. Genes upregulated in the AL treatment (downregulated in CR) are shown in green, while genes upregulated in CR (downregulated in AL) are shown in orange. The x-axis represents the number of DEGs with a significant adjusted p-value (< 0.05), corrected using the BH method. Additionally, Supplementary Figure S3 presents another bar plot depicting sex-biased gene expression in AL and CR-fed seahorses.

**Figure 4:**
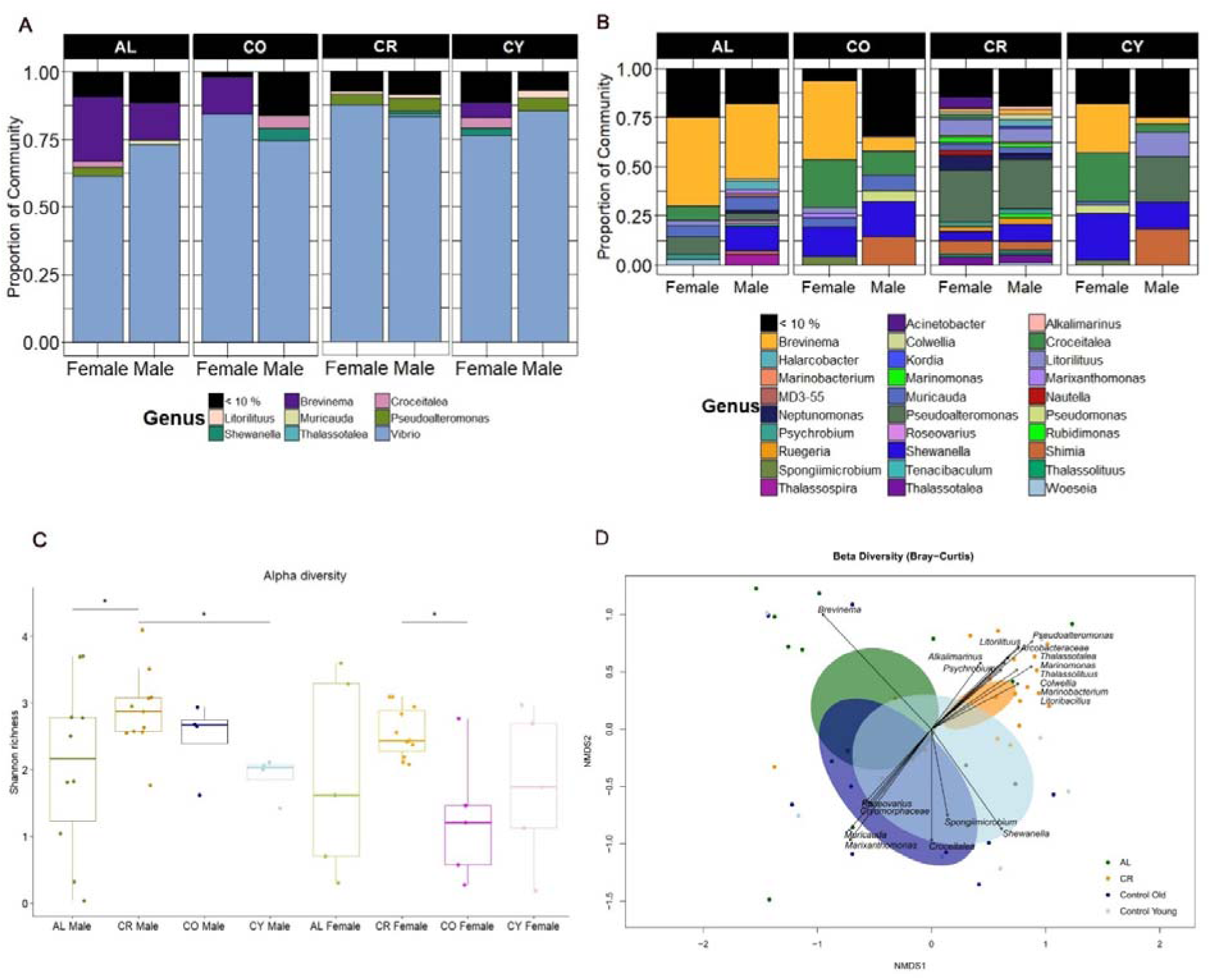
Microbial biodiversity analysis from hindgut tissue of seahorses. (A) Bar plot showing the relative abundance of microbial taxa at the genus level. Panels are categorized by diet—ad libitum (AL) and caloric restriction (CR)—as well as by age group—control old (CO) and control young (CY). Within each subcategory, data are further divided by sex (male and female). Low-abundance taxa (<10%) are grouped together. (B) Bar plot showing the relative abundance of microbial taxa at the genus level after excluding *Vibrio*, allowing for better visualization of background microbial communities. (C) Boxplots of alpha diversity (Shannon index) for diet groups (AL and CR) and age groups (CO and CY), separated by sex. P-values are derived from the post hoc Dunn test (see Table S6); significance is indicated with * as p < 0.05. (D) Non-metric multidimensional scaling (NMDS) plot based on the Bray-Curtis dissimilarity metric, comparing microbial communities across diet and age groups. Each dot represents an individual, and 95% confidence ellipses illustrate group variability based on standard error. Arrows indicate microbial taxa significantly associated with community structure (p < 0.01), highlighting key indicator species for each group.

#### 2.3.5 Correlation analysis between gut microbiota and transcriptomic data

To assess potential associations between microbiota composition and host transcriptomic profiles, we performed a Mantel test comparing the 205 bacterial genera identified in our 16S rRNA sequencing data with the normalized transcriptomic data from liver, head kidney, testes, and ovaries (Table S6). The Mantel test evaluates correlations between two distance matrices, using the Bray-Curtis dissimilarity method to assess the relationship between microbiota and gene expression patterns. Significant correlations were further explored using Multidimensional Scaling (MDS), an ordination method that reduces high-dimensional data to a lower-dimensional space for visualizing potential associations between expressed genes and bacterial taxa. To further investigate the functional implications of the significant associations (p < 0.01), we performed a gene set enrichment analysis using the g:Profiler web tool to identify enriched pathways related to the associated genes. The results of this analysis are provided in Supplementary Material, Section 3.6. Additionally, we conducted a correlation analysis specifically focused on immune-related genes and their association with bacterial taxa. The resulting associations are presented in Supplementary Figure S5.

## 3. Results

### 3.1 Effects of caloric restriction on growth and condition index

The reduced caloric intake significantly impacted growth in the CR group (Fig. 1). By the end of the experiment, the AL group had a larger overall size and was heavier compared to the CR group (Fig. 1A and S1A-D, KW-test p < 0.0001). Prior to beginning the diet treatment, the seahorses had a mean weight of 0.58 ± 0.08 g for CR and 0.50 ± 0.03 g for AL (LMM p = NS, Table S6) and a mean length of 4.5 ± 0.21 cm for CR and 4.6 ± 0.21 cm for AL (Fig. S1C, LMM p = NS, Table S6). After 157 days, the AL group had a mean weight of 8.61 ± 0.2 g, while the CR group displayed a mean weight of 3.51 ± 0.24 g (Fig. 1A, LMM p < 0.001). Across the growth trajectory monitoring we noticed already on day 55, weight measurements showing stark contrasts for male seahorses with AL 3.53 ± 0.4 g vs. CR 1.12 ± 0.07 g (LMM adj.p < 0.001, Fig. 1A) and then for both sexes on day 111 with AL male 8.2 ± 0.73 g vs. CR 3.24 ± 0.19 g (LMM adj.p < 0.0001, Fig. 1A), and AL female 4.73 ± 0.61 g vs. CR 2.5 ± 0.32 g (LMM adj.p < 0.001, Fig. 1A, Table S6).

After five months, the differences between males and females became even more pronounced. Female seahorses exhibited a slight but significant difference in weight, with AL females weighing 5.6 ± 0.9 g compared to 2.8 ± 0.33 g for CR females (Dunn-test, adj.p < 0.02, Fig. 1A and S1B). In contrast, males displayed a more substantial and highly significant weight difference, with AL males weighing 9.9 ± 0.73 g versus 4.18 ± 0.24 g for CR males (Dunn-test, adj.p < 0.0004, Fig. 1A and S1B). However, when we calculated the condition index, we observed that the CR treatment had a smaller impact on the overall condition of males (Tukey HSD, adj.p = 0.09) compared to females (Tukey HSD, adj.p < 0.002, Fig. 1B). It is important to interpret the results for female seahorses with caution, as only five AL females remained by the end of the experiment compared to 11 CR. This difference is most likely due to the initial unequal sex ratio, rather than mortality, as both male groups maintained 11 individuals each.

Throughout the dietary period, we closely monitored the fish and documented the emergence of pouch formation in male seahorses, as they were not sexually mature at the beginning of the experiment. On day 19, we observed the first signs of pouch formation in an AL, and by day 28, we were able to distinguish between males and females in all AL seahorses. On day 39, pouch formation was documented in the CR group, and by day 47, we could confidently determine the sex of all CR individuals. Ovarian weight measured at the end of the fasting experiment did not significantly differ between AL and CR groups (ANOVA, p = 0.3; Fig. 1C). This lack of significance could be attributed to the lower number of biological replicates in the AL group (n = 5) compared to the CR group (n = 11). Furthermore, at the end of the experiment (day 157), we assessed the relative male pouch size between treatment groups (Figs. 1D and S1H). AL males showed a moderate difference, with pouch size averaging approximately 19% of total body length, compared to 17% in CR males (ANOVA, p-value < 0.03, Fig. 1D).

In the supplementary materials, we also compare the condition index of the young and old control groups (Fig. S1E-G, Table S6 statistics). Specifically, we compared the condition factor between CO and CY for both males and females (Fig. S1E), and further compare the condition factor between the dietary treatment groups AL and CR with the age control groups CO and CY (Fig. S1F). Due to differences in life stages between the CR experiment and the age control groups, direct comparisons are not fully appropriate, and therefore this analysis is discussed only in the supplementary material. However, we found no significant difference in fish condition between AL and CO (Tukey HSD, adj.p = 0.9), a moderate difference between CR and CO (Tukey HSD, adj.p = 0.03) with a rather more significant effect when comparing CR and CY (Tukey HSD, adj.p = 0.007), but a very strong significant difference between AL and CY (Tukey HSD, adj.p < 0.00001, Fig. S1F).

### 3.2 Effects of caloric restriction on tissue composition

RNA sequencing (RNAseq) of the liver, head kidney, and gonads (testes and ovaries) was analysed to assess sex-specific effects of CR treatment on metabolic, immunological, and reproductive acclimatization. PERMANOVA analyses of the five-month CR and AL treatments revealed that the effects of treatment and sex varied across tissues. In the liver, both treatment and sex had significant effects (PERMANOVA; p < 0.001), but no interaction was observed (Table S6). Pairwise comparisons with Bonferroni correction identified significant differences between CR and AL males, AL females and AL males, and AL females and CR males (adj.p < 0.05). Notably, the treatment effect was weaker in females, as no significant difference was found between AL and CR females. In the head kidney, the treatment effect was not as pronounced (p < 0.04), instead, a strong sex effect was observed (p < 0.001), with the only significant interaction represented between CR females and AL males (p < 0.05). Reproductive tissues showed no strong response to CR, with no effect on testes (p = 0.6) and a non-significant trend in ovaries (p = 0.06, Table S6).

To investigate whether fasting induces physiological rejuvenation, we compared gene expression profiles of seahorses subjected to different feeding regimens (AL, CR) with those of control age cohorts (CO, CY). PERMANOVA tests revealed significant differences between dietary treatments and age groups across all tissues (p < 0.002), except for the testes (p = 0.2). In the liver, AL seahorses differed significantly from CY (adj.p = 0.006) and to a lesser extent from CO (adj.p = 0.012). Interestingly, the opposite pattern was observed for CR seahorses, with CR vs. CO (adj.p = 0.006) and CR vs. CY (adj.p = 0.012) showing reversed significance trends (Table S6). In the head kidney, AL seahorses again differed significantly from CY (adj.p = 0.006) and barely reached significance with CO (adj.p = 0.048). However, unlike the liver, CR seahorses displayed a significant difference from CY (adj.p = 0.006) but were less distinct from CO (adj.p = 0.024). As previously noted, the male testes exhibited no significant differences between treatment groups or age cohorts (Table S6). While a PERMANOVA detected an overall significant effect in the ovaries, none of the pairwise comparisons remained significant after correction.

The PCA analysis, comparing all organ types and dietary treatments, revealed that the first three principal components (PCs) were primarily driven by tissue and organ effects, without clear treatment-related differences (Figs. S2 and 2A). Specifically, PC1 separated the liver from the other organs (Fig. S2), PC2 distinguished the head kidney from the liver and gonads (Fig. 2A), and PC3 further separated male and female gonads (Fig. 2A). In contrast, PCs 4 and 5 were more influenced by dietary and age-related factors (Fig. 2B). PC4 separated the dietary groups (AL and CR) from the age cohorts (CO and CY), while PC5 distinguished CR and CY from AL and CO (Fig. 2B). These differences were most pronounced in the liver, where the separation was particularly clear (Fig. 2B).

We next explored the gene expression profiles to further investigate the effects of CR on seahorse physiology, focusing on sex differences and aging. The DGE analysis revealed considerable heterogeneity in the number of significantly differentially expressed genes across different organs (Table S1) and reflected the statistical results above. Females exhibited a stronger transcriptional response to caloric restriction (CR) in the liver, with 375 DEGs compared to 83 in males (Table S1). Sex differences were more pronounced in the AL group (501 DEGs between males and females) but were reduced under CR (83 DEGs), indicating a convergence in gene expression profiles between sexes under dietary restriction. For the age cohort comparisons, male CR induced a stronger shift in gene expression compared to AL, particularly when both treatments are compared to the same age cohort (e.g., CR vs. CO = 2454 DEGs vs. AL vs. CO = 933 DEGs). In contrast, in females, the difference between CR and AL is less pronounced, and CR appears to buffer age-related changes more effectively, as indicated by the much lower number of DEGs in CR vs. CY (601) compared to AL vs. CY (1891).

The head kidney exhibited pronounced sex differences in gene expression but showed a minimal response to caloric restriction. In females, there were no DEGs between AL and CR, indicating that dietary restriction did not significantly alter gene expression in this organ. In males, only four DEGs were detected between AL and CR, reinforcing the weak effect of caloric restriction (Table S1). However, the sex differences were striking - AL males and females differed by 973 DEGs, and this difference increased further under CR (1728 DEGs). When comparing dietary treatments across age cohorts, younger males exhibited the strongest transcriptional response in the head kidney. AL males showed almost no difference from old controls (11 DEGs) but had a substantial response when compared to young controls (1674 DEGs). Similarly, CR males showed a moderate shift compared to old controls (801 DEGs), which became much more pronounced in comparison to young controls (2846 DEGs). In females, these effects were weaker but followed a similar trend, with 502 DEGs (AL vs. old controls) and 632 DEGs (AL vs. CY), while CR resulted in 558 DEGs (vs. CO) and 1054 DEGs (vs. CY).

Although CR did not affect the male testes, the female ovaries exhibited 366 DEGs between the AL and CR treatments. When comparing the age cohorts, no significant differences were found between AL and CO or CR and CY. In contrast, there were 218 significantly DEGs between CR and CO, while a striking 4966 DEGs were found between AL and CY.

### 3.3 Gene enrichment analysis reveals sex-specific fasting effects on liver metabolism

In the female liver (AL vs. CR), the most significantly enriched pathways across GO:BP, KEGG, and REAC analyses were related to sterol and cholesterol biosynthesis (Table S2, sheet A). Genes upregulated in CR females included cytochrome P450 family 51 (*cyp51*), squalene epoxidase (*sqle*), and mevalonate kinase (*mvk*), all crucial in cholesterol and sterol biosynthesis (52). Other upregulated genes, such as hormone-sensitive lipase (*lipe*) and lipase hepatic A (*lipca*), play key roles in lipid metabolism and uptake (Table S3) (53). CR also upregulated genes involved in ATP production, oxidative stress response, and autophagy, including isocitrate dehydrogenase (*idh2*), peroxiredoxin 4 (*prdx4*), peroxiredoxin 6 (*prdx6*), autophagy-related 13 homolog (*atg13*), and acetoacetyl-CoA synthetase (*aacs*), all of which contribute to energy homeostasis and survival under nutrient depletion (54–57).

Conversely, CR downregulated genes related to steroid synthesis, including P450 family 17 subfamily A (*cyp17A2*) and the progesterone receptor (*pgr*) (58), as well as genes involved in iron metabolism and erythropoiesis, such as STEAP3 and erythropoietin receptor (*epor*) (59,60).

In males, DEGs in the liver (AL vs. CR) were primarily associated with energy production, with enriched pathways including ATP biosynthetic processes (GO:BP), oxidative phosphorylation (KEGG), and mitochondrial biogenesis (REAC) (Table S2, sheet B). Although CR affected fewer genes in males than in females, most were upregulated in fasted individuals (Fig. 3). The few downregulated genes included *coq8ab*, involved in coenzyme Q biosynthesis for mitochondrial electron transport, and *bco11*, which cleaves beta-carotene into retinal, a precursor of vitamin A (61). Some upregulated genes overlapped with those in females, such as *idh2* and *lipca* (Table S2, sheet B). However, CR males showed increased expression of *ldhd*, which converts pyruvate to lactate and regenerates NAD+, and *fetub*, a glycoprotein linked to impaired glucose metabolism and insulin resistance (62). Additional upregulated genes included *thop1*, which regulates energy metabolism, with knockout studies linking it to increased oxidative metabolism (63), and *selenop2*, a selenoprotein involved in oxidoreductase reactions (64). We also identified DEGs related to circadian rhythm regulation that were induced in expression in the liver of CR males, including clock circadian regulator b (*clockb*) and cryptochrome circadian regulator 3 (*cry3a*). The circadian rhythm is closely linked to the daily cycle of nutrient intake and metabolic control. CLOCK, a key transcription factor in the circadian clock machinery, modulates the chromatin environment by altering histone acetyltransferase (*HAT*) activity (65). *Cry3a*, an ortholog of human *cry1*, functions as a molecular clock regulator by participating in an autoregulatory transcriptional loop with an ∼24-hour periodicity, where CLOCK binds to the cry1 promoter region (66).

### 3.4 Calorie restriction reduces immune response and lipid metabolism pathways in ovaries

No significantly DEGs were found in the testes between AL and CR treatments, compared to ovaries which appeared far more affected through the diet with 356 genes being downregulated under CR and only ten upregulated (Fig. 3). Enrichment analysis highlighted key pathways, including lipid catabolic process (GO:BP), lysosome (KEGG), and innate immune system (REAC) as the most enriched (Table S2, sheet C). Among the downregulated genes in CR, several were involved in lipid catabolism, such as ATPase H+ transporting V1 and V0 (*atp6v1f, atp6v1e1b, atp6v0cb*), as well as nutrient-sensing genes that regulate longevity pathways. These included forkhead box O1a (*foxo1a*), a transcription factor that integrates insulin action with nutrient and energy homeostasis, part of the FOXO family regulating metabolism, stress resistance, apoptosis, and longevity (67). Similarly, sestrin 2 (*sesn2*), known to activate mTORC1 (68), and TELO 2 interacting protein 2 (*tti2*), which stimulates downstream cascades for cell growth via mTORC1 (69), were also downregulated. The downregulation of these genes in CR suggests a suppression of growth and cell proliferation pathways. Conversely, similar pathways were upregulated in AL ovaries, including genes like claudin 7 (*cldn7b*), involved in solute transfer across cell monolayers and possibly contributing to cancer cell survival, growth, and invasion (70), and Rhophilin Rho GTPase binding protein 2 (*rhpn2*), a scaffold protein that participates in protein complex assembly, intracellular signalling, and endosome targeting (71).

Immunological responses in the ovaries of CR females were suppressed, with multiple genes related to the innate immune system and lipid metabolism showing downregulation. Notably, two acyl-CoA synthetase (*ACS*) enzymes, ACS bubble gum (*acsbg*) and long-chain ACS (*acsl*), both involved in lipid metabolism. *Acsl*, in particular, is known to play a role in immune response in rainbow trout, being upregulated under hypoxic conditions to enhance tolerance (72,73). Alongside these lipid-metabolism-related genes, all genes involved in the innate immune pathway were also downregulated in CR ovaries (Fig. 3, Table S3 sheet D). This included genes such as cytokine-like interleukin 12 receptor beta 2a (*ilrb2a*), which stimulates interferon-γ production from T cells and NK cells (74); M17, a subfamily cytokine with similarities to mammalian IL-6 that induces macrophage activation in goldfish (75); and *mpeg1.2*, an antibacterial gene induced by infection (76). Other downregulated genes in this pathway included interferon regulatory factor 5 (*irf5*), involved in antiviral and inflammatory responses (77), chemokine receptors (*cxcr4a, cmklr1*), and tumor necrosis factor (tnf), all contributing to the inflammatory response (78). Furthermore, toll-like receptors (TLRs) 21 and 22, essential pattern-recognition receptors for detecting pathogen-associated molecular patterns (PAMPs), were also downregulated.

### 3.5 Calorie restriction promotes greater gut microbial diversity

The analysis of microbial 16S rRNAseq in seahorse hind-gut tissue found that *Vibrio* was the most abundant bacterial genus across all treatments (CR, AL), age cohorts (CO, CY), and sexes. However, its prevalence differed between treatments. In the CR group, *Vibrio* dominated the microbial community, comprising approximately 90% of the total abundance in females and 85% in males, while in the AL group, it accounted for a lower proportion, ranging from 60% in females to 75% in males (Fig. 4A). This suggests that CR creates an environment where *Vibrio* outcompetes other bacterial taxa, whereas in the AL treatment, microbial diversity appears to be higher, allowing for the presence of additional genera.

To better understand the composition of smaller bacterial communities, we reanalysed the data after excluding *Vibrio*, re-normalizing and applying CLR transformation to correct compositional biases, allowing for more accurate comparisons of less abundant taxa across all groups. This revealed a substantial presence of other taxa across all groups. In the AL treatment, *Brevinema* (family *Brevinemataceae*) emerged as the dominant genus accounting for approximately 45% of the relative abundance in females and 38% in males (Fig. 4B). In fact, β-diversity NMDS plots comparing treatment groups (AL, CR) and age cohorts (CO, CY) indicated a strong association between *Brevinema* and AL seahorses (Fig. 4D). In contrast, CR seahorses exhibited a more diverse background microbial community, though with considerably lower relative abundances of individual taxa (Fig. 4B). Among the most prominent bacterial groups in CR seahorses were several genera within the phylum Proteobacteria, including *Pseudoalteromonas, Neptunomonas, Litorilituus, Shewanella*, *Marinobacterium, Marinomonas*, and *Thalassotalea* (Figs. 4B and 4D).

Shannon diversity analysis showed no significant differences in within-individual variability between treatment groups (CR, AL) or age cohorts (CO, CY), with overall low richness values ranging from below 1 to a maximum of 2.5 (Levene’s test, p > 0.05; Fig. S4A, Table S6). This aligns with the dominance of *Vibrio* across all samples. However, when *Vibrio* was excluded, microbial diversity varied more within Al seahorses (Levene’s test, p < 0.002, Table S6), also for CO and CY females, whose Shannon richness ranged from below 1 (indicating few dominant taxa) to 3 (indicating higher diversity with several coexisting species). In contrast, most CR individuals had more consistent values around 3 (Fig. 4C). CO and CY males also exhibited higher diversity, averaging around 2–2.5. Significant differences in α-diversity were detected between several treatment groups (Table S6), including CR males vs. both AL and CY males (Dunn test, p = 0.04 and p = 0.01 respectively), and CR females vs. CO females (p = 0.02). No significant differences were detected between sexes within the same treatment group and only CO male vs. CO females barely reached significance (p = 0.053). However, significant differences were observed in comparisons where treatment, sex, and age effects were confounded, such as between CR males and CO females or CY females and CR males (Table S6).

The effect of CR on β-diversity, which measures differences in microbial community composition between groups, was the strongest driver of variation in our analysis. Significant differences were observed between CR and AL when analysing the full dataset, including *Vibrio* (adj. p < 0.006; Fig. S4C). However, this effect was even more pronounced when examining background microbial communities, excluding *Vibrio* (adj. p < 0.002; Fig. 4D). Additionally, significant differences emerged between CR and all other groups (AL, CO and CY) in the absence of *Vibrio* (adj.p < 0.002). In contrast, there was no effect of sex or sex by treatment interaction on gut microbiome composition (Fig. S4D). While no significant differences were observed in overall α- and β-diversity measures between males and females, closer observation of the relative abundance results (Fig. 4B) revealed some interesting trends in community composition across the different treatment groups for sex. For instance, *Brevinema* was similarly highly prevalent in both sexes in the AL group, compared to a lower abundance for both sexes in the CR group, while in CO and CY, females higher relative abundance compared to females. In contrast, *Croceitalea* was found to be dominant in female seahorses in the CO and CY groups, where it represented around 25% of the total microbiome composition. This dominance was notably lower in AL females (around 6%) and nearly absent in CR females. *Marinobacterium* was more abundant in CO and CY males (15%) but barely present in females, and showed low levels in both sexes in CR (ca. 5%). Finally, *Shewanella* was present in all groups and both sexes, except in AL females, where it was absent. These taxon-level shifts highlight the complex interaction between diet, sex, and microbiome composition.

## 4. Discussion

Beneficial health effects of CR are well documented across various model species (yeast, nematodes, fruit flies and rodents), with positive effects encompassing improved metabolism, reduction in inflammation to a higher diversity of the gut microbiome (79,80). This has led to the belief of evolutionarily conserved mechanisms of CR effects across the animal kingdom, which were challenged by a meta-analysis, revealing twofold stronger effects in model species than non-model species (81). This is potentially attributed to genetic sensitivity and publication bias favoring positive results in model organisms. Such bias underscores our constrained comprehension of how fasting impacts non-model systems, which exhibit different evolutionary adaptations to food availability and varied evolutionary flexibility in reproductive systems, involving female and male-specific resource allocation trade-offs (6,79). These differences could shape how males and females adjust to nutritional stress, leading to divergent fasting effects across species.

Our study aimed to elucidate the effect of CR, implemented through an alternating fasting regimen, on *Hippocampus erectus*. In this non-model species, males are the pregnant sex, incurring costly investment into development of brooding structure (82,83). Conversely, females produce energy-rich eggs. *H. erectus* unique reproductive biology allowed us to disentangle over sexual maturation, the cost of investment into the brooding structure vs. investment into egg production, two costly traits that are commonly associated with female reproduction across many taxa. Seahorses further lack a stomach leading to unknown responses to food deprivation, as they typically eat at frequent intervals (36).

The CR treatment induced a slower growth trajectory (Fig. 1A) likely through energy allocation trade-offs, which aligns with previous research, particularly when implemented during the juvenile stage (21). Compared to the AL males, the CR males exhibited reduced body mass and size after 5 months (Dunn-test, adj.p < 0.0004, Fig. 1 and S1B), this effect was slightly less in female seahorses (Dunn-test, adj.p < 0.02, Fig. 1A and S1B). Interestingly, an assessment of the condition index revealed that the CR treatment had a less pronounced impact on the overall condition of males relative to females (Tukey HSD, adj.p = 0.09 vs. adj.p < 0.002, Fig. 1B respectively). The results suggest that males are more sensitive to CR in terms of somatic growth, potentially prioritizing body maintenance over reproduction. This is reflected in the delayed pouch formation observed in CR males, which occurred approximately 20 days later than in AL males. By day 157, CR males also had a smaller relative pouch size compared to AL males (ANOVA, p < 0.03; Fig. 1D), suggesting a slower developmental trajectory. This delay may also be attributed to the fact that pouch development is highly resource-dependent, being a costly process that can only be completed when resources are available, as it is crucial for the individual’s fitness.

In contrast, females under CR maintained ovarian weight (ANOVA, p = 0.3; Fig. 1C) despite experiencing a decline in overall body condition. This suggests a trade-off between somatic maintenance and reproductive investment, with females prioritizing ovarian function even under CR. Since egg production is energetically demanding, CR females may have mobilized body reserves to sustain ovarian tissue, even at the expense of body condition. Additionally, ovarian maintenance may be less energy-intensive than brood pouch development, allowing females to preserve reproductive potential with minimal energetic input. However, it remains possible that CR negatively affected egg quality or maturation, which may not be reflected in total ovarian weight. The small sample size in the AL group (n = 5) limits our ability to detect subtle differences.

These findings suggest a sex-specific response to CR: males delay reproduction, investing in body maintenance and postponing pouch development, while females sustain reproductive investment at the cost of body condition. This aligns with the disposable soma theory, where males may defer reproduction until conditions improve, avoiding the high energetic costs of developing a vascularized pouch. Interestingly, these results contrast with patterns observed in mammals, where early-life adversity often accelerates reproductive development in males. Such differences may reflect distinct life-history strategies, with seahorse males prioritizing somatic growth to develop a larger pouch, which could support a greater number of viable offspring per clutch. Delaying pouch development to later in life may thus enhance overall fitness for males, while females allocate resources to maintaining reproductive potential despite energetic constraints (84).

The cost of nutritional egg provisioning is highlighted by the stark contrast in gene expression between males and females in both gonadal and liver tissue, despite the absence of a significant difference in female ovarian weight between CR and AL (Fig. 3). Fasting did not affect sperm production (no DEGs between AL and CR treatments in testes), consistent with findings in conventional model organisms (6), likely due to the lower production cost of sperm compared to resource-intensive egg formation. In the ovaries 356 DEGs were downregulated (opposed to 10 upregulated) upon CR (Fig. 3), with pathways relating to lipid metabolism and innate immune system (Table S2, sheet C). Also in the liver, female seahorses had more DEGs compared to males (Fig. 3) involving similar enriched pathways pertaining to metabolism of steroids (Table S2, sheet A), however, in the liver DEGs were predominantly upregulated in CR seahorses. This underscores the reliance of female reproductive functions on energy reserves in adipose tissue, where fat cells produce the hormone leptin, a process diminished during fasting with the loss of body fat (21). Decreased leptin levels correlate with lower progesterone concentrations (85), supported by downregulated *pgr* and *cyp17A2* in the liver of fasting females, which could impair oocyte maturation. Nonetheless, as mentioned earlier, females may prioritize reproduction by investing in egg production even at the cost of egg quality, as indicated by the downregulation of immune responses and lipid genes in the ovaries. This could be a strategy to produce more eggs of lower quality, rather than delaying egg production and potentially missing optimal reproductive windows due to environmental constraints.

Regardless of male seahorses showcasing only few DEGs upon CR, the liver displayed enriched pathways and genes important for aging, including genes involved in cellular respiration and oxidative phosphorylation (Table S2, sheet B) (11,21). In the liver of CR males genes controlling circadian rhythm (*cklockb* and *cry3a*), impacted by daily nutrient level fluctuations (86,87), were upregulated. Studies in mice found sex-specific signaling of circadian clock genes, with *cry1* being higher in the liver of CR males (88). A well-regulated circadian clock improves overall health, and this may be particularly important for males delaying reproduction, if relying on a longer lifespan to maximize fitness ensures they have the necessary resources for future reproductive efforts.

The downregulation of numerous inflammatory genes (including TLRs 21 and 22, chemokines, tnf, irf5, and M17, known for their pro-inflammatory roles in fish (75,77,78)) in CR female ovaries suggests a potential limitation in their capacity to prime their eggs with immunological components for transgenerational plasticity (89,90). This aligns with prior research in the syngnathid *Syngnathus typhle* (91,92) where exposure to environmental stressors, like elevated temperatures, erased the effects of transgenerational immune priming, potentially driven by a resource allocation trade-off between survival and immune priming in times of limited resources.

We observed potential beneficial effects of fasting by upregulated genes for increased autophagy (*atg13*) and ketosis (*aacs*) in livers of CR seahorses, in line with other studies (21). The upregulation of both *sesn2* and *tti2* in AL compared to CR (ovarian tissue) results in an activation of mTORC1 and cell growth (68,69). Suppressing mTOR activity may promote longevity by shifting the body from a state of growth to autophagy, thus improving cell homeostasis (11,21). Additionally, *foxO1a* was upregulated in AL ovaries (Fig. 3), *FoxO* is found typically elevated in response to high levels of reactive oxygen species (ROS) having a dual role in combating oxidative stress, being both pro-survival and pro-apoptotic (93). *FoxO1a* can have positive effects by promoting DNA repair (93). A potential rejuvenation effect in the ovaries of CR females was revealed by higher lipid stores and increased lipid synthesis in AL female ovaries promoting cell growth compared to fasted females prioritized pathways for repair and regeneration, shifting from high reproductive output to increased somatic maintenance (Tables S1 and S2, sheet D). Despite the absence of notable morphological effects of CR in females, lipid metabolic pathways were upregulated in the liver and downregulated in the ovaries.

The gut microbiome is crucial for various aspects of health, including vitamin synthesis, immune system fine-tuning, maintaining the integrity of the intestinal barrier, and the modification of bile acids, neurotransmitters, and hormones (94,95). Fasting induces changes in mucus production, which in turn affects gut microbial diversity (96). The gut microbiota play a pivotal role in influencing fish health by fostering the development of the gut epithelium, enhancing the immune system’s functioning and acting as protective barrier, impeding the colonization of harmful pathogens within the gut (97,98). In the CR treatment, *Vibrio* dominated the microbial community, comprising approximately 90% of the abundance in females and 85% in males, significantly higher than in the AL treatment (60% in females and 75% in males) (Fig. 4A). This suggests that CR conditions favor *Vibrio* dominance, potentially by creating a more competitive environment where *Vibrio* outcompetes other taxa under resource-limited conditions. *Vibrio* species can have both beneficial and pathogenic effects, depending on environmental factors and host interactions, which may influence the overall health and microbiome composition of the seahorses (99).

When *Vibrio* was excluded from the analysis, the CR group showed a more diverse microbial background, with several Proteobacteria genera such as *Pseudoalteromonas, Neptunomonas, Litorilituus*, and others emerging as prominent taxa (Fig.4B). CR seahorses were the strongest driver of variation in the beta-diversity analysis, with CR individuals being significantly distinct to all other groups (adj.p < 0.002, Fig. 4D), further emphasizing that CR profoundly reshapes the microbial community. The heightened relative abundance and taxonomic diversity within the *Proteobacteria* phylum (Figs. S4B), aligns with previous studies suggesting an increased relative abundance of *Proteobacteria* in fasting vertebrates, including fish (98,100). *Proteobacteria*, recognized for their crucial role in preparing the gut for colonization by strict anaerobic bacteria, play a vital part in maintaining gut health (101). In contrast, the AL group exhibited a more balanced microbial community, where the genus *Brevinema* (phylum *Spirochaetota*), emerged as the dominant non-*Vibrio* genus (Fig. 4B).

*Brevinema* relative abundance was closely associated to the expression of several immune genes in the testes (e.g., irf10, mhc1zca, cd74; Figure S5). This association seems to be particularly relevant in the AL treatment, as *Brevinema* was severely reduced in the gut of CR individuals compared to AL (Fig. 4B). *Brevinema* is not only a key player in microbial communities associated with fish mucus (102) and gut microbiota (103), but was also identified as a microbe that is involved in paternal-specific vertical microbial transfer (104). It is tempting to develop the hypothesis that CR reduced relative abundance of *Brevinema* connected to a lowered expression of several immune genes results in reduced trans-generational microbial and immune transfer also on the paternal side under stressful environmental conditions (92).While testing this hypothesis will require investigations on organ-specific microbiomes, we can currently state more general that *Brevinema* may play a critical role in immune function, with CR potentially disrupting this relationship and leading to diminished immune responses.

Although beta diversity analysis revealed no significant differences between sexes and treatments (Fig. S4D), indicating that the major driver of microbial community differences is the dietary treatment, significant differences in alpha diversity were observed. Specifically, alpha diversity was higher in CR individuals, with a notable difference between CR and AL males (Dunn test, p = 0.04), but not females, suggesting that fasting influenced microbial evenness in a sex-dependent manner. Most types of fasting were shown to increase α-diversity (79), and our findings align with this background pattern.

On the genus level, *Pseudoalteromonas* exhibited the highest relative abundance (approximately 25% in both males and females) compared to AL, comprising marine species with diverse activities including anti-bacterial, algicidal, anti-viral, and specific isolates hindering the settlement of common fouling organisms (105). This could confer significant benefits to the immunological health of fasted seahorses. Among the other prevalent taxa in CR, many are commonly found in marine organisms, but their correlation with specific metabolic roles in animals remains unexplored. However, we did find *Neptunomonas*, which has potential nitrate reduction properties (106), and *Thalassotalea*, characterized as an aerobic, chemo-organotrophic genus of bacteria involved in nutrient cycling and the decomposition of organic matter (107). The efficient utilization of various forms of energy from the environment by these microbes may contribute to their abundance, providing energy for the host during fasting. Additionally, a study revealed that *Litorilituus sediminis* was associated with tumor suppressive activity (108), that could also confer some protective properties to CR seahorses. In summary, while the gut in CR conditions may exhibit potential for improved health compared to AL, a clear rejuvenation was not evident. The dominance of certain taxa in CO and CY, consistent across age groups with some variation between sexes, prompts us to consider that a younger gut does not necessarily equate to a healthier state in seahorses.

In conclusion, our findings shed light on the intricate sex-specific responses to fasting in seahorses, providing valuable insights into the adaptive strategies employed by males and females to cope with nutritional stress and maintain reproductive success. Our study emphasizes how the primary determinants of resource allocation are closely tied to sex-specific life-history strategies. In *H. erectus* males, this is exemplified by the unique focus on male pregnancy and the allocation of resources post-copulation, contrasting with the pre-copulatory investment in female eggs, leading to distinct morphological and regulatory effects of fasting between males and females, that are opposed to effects in typical model organism. Nonetheless, conserved mechanisms of CR were evident in both sexes, but to different degrees, with increased autophagy and ketosis pathways in CR females, and enrichment of pathways and genes important for cellular respiration, and circadian rhythm control in CR males. As we move forward, additional research in a diverse range of model species with varying reproductive strategies and sex roles will be essential to build upon our findings. Understanding the intricate connections between reproduction, resource availability, and fasting effects can contribute to a more comprehensive understanding of sex-specific health, lifespan dimorphism, and evolutionary adaptations in response to nutrient scarcity, particularly in the context of climate change. In an ecological context, these sex-specific fasting responses could have important implications for species survival, particularly in environments where nutrient availability fluctuates, influencing reproductive timing and success, and shaping evolutionary strategies in the face of climate change. By adopting a multidisciplinary approach, researchers can make significant strides in translating the findings into practical applications for health and beyond.

## Acknowledgments

We gratefully acknowledge Silke-Mareike Marten for her valuable input as a lab technician and our animal-care technicians Fabian Wendt and Johannes Hasse for providing support with the fish care and aquaria maintenance. Financial support was provided by the Deutsche Forschungsgemeinschaft (DFG, German Research Foundation) through the Research Training Group for Translational Evolutionary Research (RTG 2501 TransEvo) and through the CRC 1182 origin and function of metaorganismus for the possibility to genotype the microbiota.

## Data availability statement

Additional information is available in the supplementary materials. Refer to Tables S1 to S6 for comprehensive details: Table S1 presents an overview of numerical outcomes derived from the *limma* DGE analysis, encompassing various groups and organs. For the results of the GO analysis, consult Table S2. Visual insights from the DGE analysis concerning the specified organs and treatment groups are presented in Figure S3. Table S4 offers the results of the 16S rRNA amplicon sequencing analysis conducted in QIIME2, detailing all OTUs at the genus level. Table S5 encompasses morphological raw data, specifically the length and weight measurements of all CR, AL, CY, and CO seahorses. Table S6 displays the full statistical analysis results for morphology, RNA-Seq and 16S rRNA-Seq. The raw sequencing RNA-Seq data, 16S rRNA-Seq and metadata used in this study are available from the National Center for Biotechnology Information (NCBI) Sequence Read Archive (SRA) under BioProject ID PRJNA1004169 (RNA-Seq: SUB13725814; 16S rRNA-Seq: SUB13759493).

## Ethics statement

The work was carried out in accordance with the German animal welfare law and with the ethical approval given by the Schleswig-Holstein Ministerium für Energiewende, Landwirtschaft, Umwelt, Natur und Ditgitalisierung (MELUND) (permit no. 1315/2021). No wild endangered species were used in this investigation.

## References

1. Trivers RL. Parental Investment and Sexual Selection., In B. Campbell (Ed.). Sexual selection and the descent of man. 1972;

2. Hayward A, Gillooly JF. The Cost of Sex: Quantifying Energetic Investment in Gamete Production by Males and Females. Voolstra C, editor. PLoS ONE. 2011 Jan 24;6(1):e16557.

3. Speakman JR. The physiological costs of reproduction in small mammals. Phil Trans R Soc B. 2008 Jan 27;363(1490):375–98.

4. Clutton-Brock TH, Parker GA. Potential Reproductive Rates and the Operation of Sexual Selection. The Quarterly Review of Biology. 1992 Dec;67(4):437–56.

5. Janicke T, Häderer IK, Lajeunesse MJ, Anthes N. Evolutionary Biology: Darwinian sex roles confirmed across the animal kingdom. Science Advances. 2016;2(2):1–11.

6. Kane AE, Sinclair DA, Mitchell JR, Mitchell SJ. Sex differences in the response to dietary restriction in rodents. Current Opinion in Physiology. 2018 Dec;6:28–34.

7. Regan JC, Khericha M, Dobson AJ, Bolukbasi E, Rattanavirotkul N, Partridge L. Sex difference in pathology of the ageing gut mediates the greater response of female lifespan to dietary restriction. eLife. 2016 Feb 16;5:e10956.

8. McKay A, Costa EK, Chen J, Hu CK, Chen X, Bedbrook CN, et al. An automated feeding system for the African killifish reveals the impact of diet on lifespan and allows scalable assessment of associative learning. eLife. 2022 Nov 10;11:e69008.

9. Zucker I, Prendergast BJ, Beery AK. Pervasive Neglect of Sex Differences in Biomedical Research. Cold Spring Harb Perspect Biol. 2021 Oct 14;a039156.

10. Pasin C, Consiglio CR, Huisman JS, De Lange AMG, Peckham H, Vallejo-Yagüe E, et al. Sex and gender in infection and immunity: addressing the bottlenecks from basic science to public health and clinical applications. R Soc open sci. 2023 Jul;10(7):221628.

11. Sinclair DA, Howitz KT. Dietary Restriction, Hormesis, and Small Molecule Mimetics. In: Handbook of the Biology of Aging. 2005.

12. Kirkwood TBL, Shanley DP. Food restriction, evolution and ageing. Mechanisms of Ageing and Development. 2005 Sep;126(9):1011–6.

13. Masoro EJ, Austad SN. The Evolution of the Antiaging Action of Dietary Restriction: A Hypothesis. The Journals of Gerontology Series A: Biological Sciences and Medical Sciences. 1996 Nov 1;51A(6):B387–91.

14. Shanley DP, Kirkwood TBL. Caloric restriction does not enhance longevity in all species and is unlikely to do so in humans. Biogerontology. 2006 Jun;7(3):165–8.

15. Ikeya K, Kume M. Seasonal Feeding Rhythm Associated with Fasting Period of *Pangasianodon gigas*l: Long-Term Monitoring in an Aquarium. Zoological Science. 2011 Aug;28(8):545–9.

16. Wang T, Hung CCY, Randall DJ. The comparative physiology of food deprivation: from feast to famine. Annu Rev Physiol. 2006 Jan 1;68(1):223–51.

17. Keller L. Queen lifespan and colony characteristics in ants and termites. Insectes Sociaux. 1998 Aug 1;45(3):235–46.

18. Remolina SC, Hughes KA. Evolution and mechanisms of long life and high fertility in queen honey bees. AGE. 2008 Sep;30(2–3):177–85.

19. Dirks AJ, Leeuwenburgh C. Caloric restriction in humansl: Potential pitfalls and health concerns. Mechanisms of Ageing and Development. 2006;127:1–7.

20. Liao CY, Rikke BA, Johnson TE, Diaz V, Nelson JF. Genetic variation in the murine lifespan response to dietary restriction: from life extension to life shortening. Aging Cell. 2010 Feb;9(1):92–5.

21. Speakman JR, Mitchell SE. Caloric restriction. Molecular Aspects of Medicine. 2011;32(3):159–221.

22. Mitchell SJ, Scheibye-Knudsen M, Longo DL, De Cabo R. Animal Models of Aging Research: Implications for Human Aging and Age-Related Diseases. Annu Rev Anim Biosci. 2015 Feb 16;3(1):283–303.

23. Benvenuto C, Coscia I, Chopelet J, Sala-Bozano M, Mariani S. Ecological and evolutionary consequences of alternative sex-change pathways in fish. Scientific Reports. 2017;7(1):1–12.

24. Smith C, Wootton RJ. The costs of parental care in teleost fishes. Reviews in Fish Biology and Fisheries. 1995;5(1):7–22.

25. Balasubramanian P, Howell PR, Anderson RM. Aging and Caloric Restriction Research: A Biological Perspective With Translational Potential. EBioMedicine. 2017 Jul;21:37–44.

26. Stölting KN, Wilson AB. Male pregnancy in seahorses and pipefish: Beyond the mammalian model. BioEssays. 2007;29(9):884–96.

27. Lin Q, Li G, Qin G, Lin J, Huang L, Sun H, et al. The dynamics of reproductive rate, offspring survivorship and growth in the lined seahorse, Hippocampus erectus Perry, 1810. Biology Open. 2012 Apr 15;1(4):391–6.

28. López-Otín C, Blasco MA, Partridge L, Serrano M, Kroemer G. Hallmarks of aging: An expanding universe. Cell. 2023 Jan;186(2):243–78.

29. Mercken EM, Carboneau BA, Krzysik-Walker SM, De Cabo R. Of mice and men: The benefits of caloric restriction, exercise, and mimetics. Ageing Research Reviews. 2012 Jul;11(3):390–8.

30. Parrella E, Longo VD. The chronological life span of Saccharomyces cerevisiae to study mitochondrial dysfunction and disease. Methods. 2008 Dec;46(4):256–62.

31. Braeckman BP, Demetrius L, Vanfleteren JR. The dietary restriction effect in C. elegans and humans: is the worm a one-millimeter human? Biogerontology. 2006 Jun;7(3):127–33.

32. Burger JMS, Buechel SD, Kawecki TJ. Dietary restriction affects lifespan but not cognitive aging in Drosophila melanogaster: Dietary restriction and memory. Aging Cell. 2010 Feb 12;9(3):327–35.

33. Selman C, Lingard S, Choudhury AI, Batterham RL, Claret M, Clements M, et al. Evidence for lifespan extension and delayed age–related biomarkers in insulin receptor substrate 1 null mice. FASEB j. 2008 Mar;22(3):807–18.

34. Maegawa S, Lu Y, Tahara T, Lee JT, Madzo J, Liang S, et al. Caloric restriction delays age-related methylation drift. Nat Commun. 2017 Sep 14;8(1):539.

35. Zeng T, Cui H, Tang D, Garside GB, Wang Y, Wu J, et al. Short-term dietary restriction in old mice rejuvenates the aging-induced structural imbalance of gut microbiota. Biogerontology. 2019 Dec;20(6):837–48.

36. Woods CMC. Natural diet of the seahorse *Hippocampus abdominalis*. New Zealand Journal of Marine and Freshwater Research. 2002 Sep;36(3):655–60.

37. Bhatwalkar SB, Mondal R, Krishna SBN, Adam JK, Govender P, Anupam R. Antibacterial Properties of Organosulfur Compounds of Garlic (Allium sativum). Front Microbiol. 2021 Jul 27;12:613077.

38. Korsch M, Marten S, Walther W, Vital M, Pieper DH, Dötsch A. Impact of dental cement on the perilimplant biofilmlmicrobial comparison of two different cements in an in vivo observational study. Clin Implant Dent Rel Res. 2018 Oct;20(5):806–13.

39. R Core Team. R: A language and environment for statistical computing. [Internet]. Vienna, Austria: R Foundation for Statistical Computing; 2022. Available from: https://www.r-project.org/

40. Robinson ML, Gomez-Raya L, Rauw WM, Peacock MM. Fulton’s body condition factor K correlates with survival time in a thermal challenge experiment in juvenile Lahontan cutthroat trout (Oncorhynchus clarki henshawi). Journal of Thermal Biology. 2008 Aug;33(6):363–8.

41. Andrews S. http://www.bioinformatics.babraham.ac.uk/projects/fastqc. 2010. FastQC: a quality control tool for high throughput sequence data.

42. Dobin A, Davis CA, Schlesinger F, Drenkow J, Zaleski C, Jha S, et al. STAR: ultrafast universal RNA-seq aligner. Bioinformatics. 2013 Jan 1;29(1):15–21.

43. Alvarez RV, Pongor LS, Mariño-Ramírez L, Landsman D. TPMCalculator: One-step software to quantify mRNA abundance of genomic features. Bioinformatics. 2019 Jun 1;35(11):1960–2.

44. Robinson MD, McCarthy DJ, Smyth GK. edgeRl: a Bioconductor package for differential expression analysis of digital gene expression data. Bioinformatics. 2010 Jan 1;26(1):139–40.

45. Ritchie ME, Phipson B, Wu D, Hu Y, Law CW, Shi W, et al. Limma powers differential expression analyses for RNA-sequencing and microarray studies. Nucleic Acids Research. 2015;43(7):e47.

46. Law CW, Chen Y, Shi W, Smyth GK. voom: precision weights unlock linear model analysis tools for RNA-seq read counts. Genome Biol. 2014;15(2):R29.

47. Bolyen E, Rideout JR, Dillon MR, Bokulich NA, Abnet CC, Al-Ghalith GA, et al. Reproducible, interactive, scalable and extensible microbiome data science using QIIME 2. Nat Biotechnol. 2019 Aug;37(8):852–7.

48. Price MN, Dehal PS, Arkin AP. FastTree: Computing Large Minimum Evolution Trees with Profiles instead of a Distance Matrix. Molecular Biology and Evolution. 2009 Jul 1;26(7):1641– 50.

49. Yilmaz P, Parfrey LW, Yarza P, Gerken J, Pruesse E, Quast C, et al. The SILVA and “All-species Living Tree Project (LTP)” taxonomic frameworks. Nucl Acids Res. 2014 Jan;42(D1):D643–8.

50. Oksanen J, Simpson GL, F. Guillaume Blanchet FG, Roeland Kindt, Pierre Legendre, Peter R. Minchin. Community Ecology Package. 2020;

51. Arbizu MP. Arbizu MP. pairwiseAdonis: Pairwise multilevel comparison using adonis. R package version 0.4. 2020;

52. Cirmena G, Franceschelli P, Isnaldi E, Ferrando L, De Mariano M, Ballestrero A, et al. Squalene epoxidase as a promising metabolic target in cancer treatment. Cancer Letters. 2018 Jul;425:13–20.

53. Santamarina-Fojo S, González-Navarro H, Freeman L, Wagner E, Nong Z. Hepatic Lipase, Lipoprotein Metabolism, and Atherogenesis. ATVB. 2004 Oct;24(10):1750–4.

54. Reitman ZJ, Yan H. Isocitrate Dehydrogenase 1 and 2 Mutations in Cancer: Alterations at a Crossroads of Cellular Metabolism. JNCI Journal of the National Cancer Institute. 2010 Jul 7;102(13):932–41.

55. Sue Goo Rhee. Overview on Peroxiredoxin. Molecules and Cells. 2016 Jan 31;39(1):1–5.

56. Bergstrom JD. The lipogenic enzyme acetoacetyl-CoA synthetase and ketone body utilization for denovo lipid synthesis, a review. Journal of Lipid Research. 2023 Aug;64(8):100407.

57. Tian E, Wang F, Han J, Zhang H. epg-functions in autophagy-regulated processes and may encode a highly divergent Atg13 homolog in C. elegans. Autophagy. 2009 Jul;5(5):608–15.

58. Uno T, Ishizuka M, Itakura T. Cytochrome P450 (CYP) in fish. Environmental Toxicology and Pharmacology. 2012 Jul;34(1):1–13.

59. Ohgami RS, Campagna DR, McDonald A, Fleming MD. The Steap proteins are metalloreductases. Blood. 2006 Aug 15;108(4):1388–94.

60. Rossert J, Eckardt KU. Erythropoietin receptors: their role beyond erythropoiesis. Nephrology Dialysis Transplantation. 2005 Jun 1;20(6):1025–8.

61. Helgeland H, Sodeland M, Zoric N, Torgersen JS, Grammes F, Von Lintig J, et al. Genomic and functional gene studies suggest a key role of beta-carotene oxygenase 1 like (bco1l) gene in salmon flesh color. Sci Rep. 2019 Dec 27;9(1):20061.

62. Meex RC, Hoy AJ, Morris A, Brown RD, Lo JCY, Burke M, et al. Fetuin B Is a Secreted Hepatocyte Factor Linking Steatosis to Impaired Glucose Metabolism. Cell Metabolism. 2015 Dec;22(6):1078–89.

63. Ferro ES, Gewehr MCF, Navon A. Thimet Oligopeptidase Biochemical and Biological Significances: Past, Present, and Future Directions. Biomolecules. 2020 Aug 24;10(9):1229.

64. Lei XG, Combs GF, Sunde RA, Caton JS, Arthington JD, Vatamaniuk MZ. Dietary Selenium Across Species. Annu Rev Nutr. 2022 Aug 22;42(1):337–75.

65. Doi M, Hirayama J, Sassone-Corsi P. Circadian Regulator CLOCK Is a Histone Acetyltransferase. Cell. 2006 May;125(3):497–508.

66. Sancar A. Regulation of the Mammalian Circadian Clock by Cryptochrome. Journal of Biological Chemistry. 2004 Aug;279(33):34079–82.

67. Cheng Z, White MF. Targeting Forkhead Box O1 from the Concept to Metabolic Diseases: Lessons from Mouse Models. Antioxidants & Redox Signaling. 2011 Feb 15;14(4):649–61.

68. Wolfson RL, Chantranupong L, Saxton RA, Shen K, Scaria SM, Cantor JR, et al. Sestrin2 is a leucine sensor for the mTORC1 pathway. Science. 2016 Jan;351(6268):43–8.

69. Brown MC, Gromeier M. MNK inversely regulates TELO2 vs. DEPTOR to control mTORC1 signaling. Molecular & Cellular Oncology. 2017 May 4;4(3):e1306010.

70. Dahiya N, Becker KG, Wood, WH, Zhang Y, Morin PJ. Claudin-7 Is Frequently Overexpressed in Ovarian Cancer and Promotes Invasion. Campbell M, editor. PLoS ONE. 2011 Jul 15;6(7):e22119.

71. Steuve S, Devosse T, Lauwers E, Vanderwinden JM, André B, Courtoy PJ, et al. Rhophilin-2 is targeted to late-endosomal structures of the vesicular machinery in the presence of activated RhoB. Experimental Cell Research. 2006 Dec;312(20):3981–9.

72. Lopes-Marques M, Machado AM, Ruivo R, Fonseca E, Carvalho E, Castro LFC. Expansion, retention and loss in the Acyl-CoA synthetase “Bubblegum” (Acsbg) gene family in vertebrate history. Gene. 2018 Jul;664:111–8.

73. Ma F, Zou Y, Ma L, Ma R, Chen X. Evolution, characterization, and immune response function of long-chain acyl-CoA synthetase genes in rainbow trout (Oncorhynchus mykiss) under hypoxic stress. Comparative Biochemistry and Physiology Part B: Biochemistry and Molecular Biology. 2022 Jun;260:110737.

74. Fujiwara A, Suetake H, Suzuki Y, Nakanishi T, Ototake M, Yoshiura Y, et al. Identification and characterization of Fugu orthologues of mammalian interleukin-12 subunits. Immunogenetics. 2003 Aug 1;55(5):296–306.

75. Hanington PC, Belosevic M. Interleukin-6 family cytokine M17 induces differentiation and nitric oxide response of goldfish (Carassius auratus L.) macrophages. Developmental & Comparative Immunology. 2007;31(8):817–29.

76. Benard EL, Racz PI, Rougeot J, Nezhinsky AE, Verbeek FJ, Spaink HP, et al. Macrophage-Expressed Perforins Mpeg1 and Mpeg1.2 Have an Anti-Bacterial Function in Zebrafish. J Innate Immun. 2015;7(2):136–52.

77. Xia J, Hu GB, Dong XZ, Liu QM, Zhang SC. Molecular characterization and expression analysis of interferon regulatory factor 5 (IRF-5) in turbot, Scophthalmus maximus. Fish & Shellfish Immunology. 2012 Jan;32(1):211–8.

78. Mokhtar DM, Zaccone G, Alesci A, Kuciel M, Hussein MT, Sayed RKA. Main Components of Fish Immunity: An Overview of the Fish Immune System. Fishes. 2023 Feb 5;8(2):93.

79. Moatt JP, Nakagawa S, Lagisz M, Walling CA. The effect of dietary restriction on reproduction: a meta-analytic perspective. BMC Evol Biol. 2016 Dec;16(1):199.

80. Sinclair DA. Toward a unified theory of caloric restriction and longevity regulation. Mechanisms of Ageing and Development. 2005 Sep;126(9):987–1002.

81. Nakagawa S, Lagisz M, Hector KL, Spencer HG. Comparative and meta-analytic insights into life extension via dietary restriction: Dietary restriction and longevity: meta-analysis. Aging Cell. 2012 Jun;11(3):401–9.

82. Wilson AB, Orr JW. The evolutionary origins of Syngnathidae: Pipefishes and seahorses. Journal of Fish Biology. 2011;

83. Nikolas Stölting K. Male Pregnancy in the Seahorse (Hippocampus abdominalis): Investigating the Genetic Regulation of a Complex Reproductive Trait. 2010.

84. Isaac JL. Potential causes and life-history consequences of sexual size dimorphism in mammals. Mammal Review. 2005 Jan;35(1):101–15.

85. Pérez-Pérez A, Sánchez-Jiménez F, Maymó J, Dueñas JL, Varone C, Sánchez-Margalet V. Role of leptin in female reproduction. Clinical Chemistry and Laboratory Medicine (CCLM) [Internet]. 2015 Jan 1 [cited 2023 Aug 3];53(1). Available from: https://www.degruyter.com/document/doi/10.1515/cclm-2014-0387/html

86. Longo VD, Panda S. Fasting, Circadian Rhythms, and Time-Restricted Feeding in Healthy Lifespan. Cell Metabolism. 2016 Jun;23(6):1048–59.

87. Manoogian ENC, Chow LS, Taub PR, Laferrère B, Panda S. Time-restricted Eating for the Prevention and Management of Metabolic Diseases. Endocrine Reviews. 2022 Mar 9;43(2):405–36.

88. Astafev AA, Patel SA, Kondratov RV. Calorie restriction effects on circadian rhythms in gene expression are sex dependent. Sci Rep. 2017 Aug 29;7(1):9716.

89. Beemelmanns A, Roth O. Biparental immune priming in the pipefish Syngnathus typhle. Zoology. 2016 Aug 1;119(4):262–72.

90. Roth O, Klein V, Beemelmanns A, Scharsack JP, Reusch TBH. Male pregnancy and biparental immune priming. American Naturalist. 2012 Dec;180(6):802–14.

91. Goehlich H, Sartoris L, Wagner KS, Wendling CC, Roth O. Pipefish Locally Adapted to Low Salinity in the Baltic Sea Retain Phenotypic Plasticity to Cope With Ancestral Salinity Levels. Front Ecol Evol. 2021 Mar 25;9:626442.

92. Roth O, Landis SH. Trans-generational plasticity in response to immune challenge is constrained by heat stress. Evol Appl. 2017 Jun;10(5):514–28.

93. Klotz LO, Sánchez-Ramos C, Prieto-Arroyo I, Urbánek P, Steinbrenner H, Monsalve M. Redox regulation of FoxO transcription factors. Redox Biology. 2015 Dec;6:51–72.

94. Zheng X, Wang S, Jia W. Calorie restriction and its impact on gut microbial composition and global metabolism. Frontiers of Medicine. 2018;12(6):634–44.

95. Doms S, Hermes BM, Baines FJ. Evolutionary perspectives on the human gut microbiome. In: Haller D, editor. The Gut Microbiome in Health and Disease. Springer International Publishing; 2018. p. 67–78.

96. Angoorani P, Ejtahed HS, Hasani-Ranjbar S, Siadat SD, Soroush AR, Larijani B. Gut microbiota modulation as a possible mediating mechanism for fasting-induced alleviation of metabolic complications: a systematic review. Nutr Metab (Lond). 2021 Dec 14;18(1):105.

97. Roeselers G, Mittge EK, Stephens WZ, Parichy DM, Cavanaugh CM, Guillemin K, et al. Evidence for a core gut microbiota in the zebrafish. ISME J. 2011 Oct;5(10):1595–608.

98. Li T, Qi M, Gatesoupe FJ, Tian D, Jin W, Li J, et al. Adaptation to Fasting in Crucian Carp (Carassius auratus): Gut Microbiota and Its Correlative Relationship with Immune Function. Microb Ecol. 2019 Jul;78(1):6–19.

99. Baker-Austin C, Oliver JD, Alam M, Ali A, Waldor MK, Qadri F, et al. Vibrio spp. infections. Nat Rev Dis Primers. 2018 Jun 21;4(1):1–19.

100. McCue MD, Passement CA, Meyerholz DK. Maintenance of Distal Intestinal Structure in the Face of Prolonged Fasting: A Comparative Examination of Species From Five Vertebrate Classes. Anat Rec. 2017 Dec;300(12):2208–19.

101. Shin NR, Whon TW, Bae JW. Proteobacteria: microbial signature of dysbiosis in gut microbiota. Trends in Biotechnology. 2015 Sep;33(9):496–503.

102. Li T, Long M, Gatesoupe FJ, Zhang Q, Li A, Gong X. Comparative Analysis of the Intestinal Bacterial Communities in Different Species of Carp by Pyrosequencing. Microb Ecol. 2015 Jan;69(1):25–36.

103. Lyons PP, Turnbull JF, Dawson KA, Crumlish M. Effects of lowllevel dietary microalgae supplementation on the distal intestinal microbiome of farmed rainbow trout *Oncorhynchus mykiss* (Walbaum). Aquac Res. 2017 May;48(5):2438–52.

104. Beemelmanns A, Poirier M, Bayer T, Kuenzel S, Roth O. Microbial embryonal colonization during pipefish male pregnancy. Sci Rep. 2019 Jan 9;9(1):3.

105. Holmström C, Kjelleberg S. Marine Pseudoalteromonas species are associated with higher organisms and produce biologically active extracellular agents. FEMS Microbiology Ecology. 1999 Dec;30(4):285–93.

106. Liu A, Zhang XY, Chen CX, Xie BB, Qin QL, Liu C, et al. Neptunomonas qingdaonensis sp. nov., isolated from intertidal sand. International Journal of Systematic and Evolutionary Microbiology. 2013 May 1;63(Pt_5):1673–7.

107. Zhang Y, Tang K, Shi X, Zhang XH. Description of Thalassotalea piscium gen. nov., sp. nov., isolated from flounder (Paralichthys olivaceus), reclassification of four species of the genus Thalassomonas as members of the genus Thalassotalea gen. nov. and emended description of the genus Thalassomonas. International Journal of Systematic and Evolutionary Microbiology. 2014 Apr 1;64(Pt_4):1223–8.

108. Wang Y, Zhao R, Ji S, Li Z, Yu T, Li B, et al. Litorilituus sediminis gen. nov. sp. nov., isolated from coastal sediment of an amphioxus breeding zone in Qingdao, China. Antonie van Leeuwenhoek. 2013 Sep;104(3):423–30.

