## Supplementary material for "Surviving on limited resources: effects of caloric restriction on growth, gene expression and gut microbiota in a species with male pregnancy (*Hippocampus erectus*)": Table S1 and supplementary figures

**3. Results**

**3.1 Effects of caloric restriction on growth and condition index**

**Figure S1: Morphological and physiological differences across dietary treatments and age**

**Figure A:** This boxplot illustrates the comparison of seahorse weight (g) between the ad libitum (AL) and calorie-restricted (CR) dietary treatments over the course of 157 days. Prior to the dietary treatment, the seahorses exhibited a mean weight of 0.58 ± 0.08 g for CR and 0.50 ± 0.03 g for AL (LMM p.val = NS) After 157 days, the AL group demonstrated a mean weight of 8.61 ± 0.2 g, while the CR group displayed a mean weight of 3.51 ± 0.24 g.

**Figure B:** This boxplot shows weight differences for 11 AL males, 11 CR males, 5 AL females, and 11 CR females on the last day of treatment (day 157). Females showed a slight significant difference in weight (AL 5.6 ± 0.9 g vs. CR 2.8 ± 0.33 g, Dunn-test adj.p.val < 0.02, Fig. S1B), but males showed a strong significant difference in weight (AL 9.9 ± 0.73 g vs. CR 4.18 ± 0.24 g, Dunn-test adj.p.val < 0.0004, Fig. S1B).

**Figure C:** This boxplot focuses on the length differences between the AL and CR dietary treatments. The AL group had a mean length of 4.5 ± 0.21 cm for CR and 4.6 ± 0.21 cm for AL on day 1, and on day 157 12.93 ± 0.42 cm for AL, while the CR group had a mean length of 10.28 ± 0.23 cm.

**Figure D:** In this boxplot, we analyse the size differences between the dietary treatments while considering the sex of the seahorses. For males, the AL group (13.75 ± 0.38cm) displayed significantly greater length than the CR group (10.76 ± 0.11 cm, Tukey HSD adj.p < 0.0001). In contrast, for females, there was no significant difference in length between the AL (11.11 ± 0.35 cm) and CR (9.81 ± 0.41 cm) groups.

**Figure E:** This figure compares the condition index (Fulton’s condition factor: weight/length³) between young (CY) and old (CO) seahorses, separated by sex. The x-axis represents the four groups based on age and sex, while the y-axis shows Fulton’s condition factor. There was a significant difference in condition index between old and young seahorses. In males, the difference between CO and CY groups was marginally significant (Tukey HSD adj. p = 0.05), whereas in females, the CO vs. CY comparison was more statistically significant (Tukey HSD adj. p = 0.03). For reference, the average weight and length of each group were as follows:

CO males: 12.16 ± 1.5 g, 15.12 ± 0.4 cm ; CO females: 8.13 ± 0.6 g, 12.96 ± 0.3 cm CY males: 1.13 ± 0.07 g, 7.88 ± 0.2 cm; CY females: 1.23 ± 0.19 g, 7.90 ± 0.2 cm.

**Figure F:** This figure compares the condition index (Fulton’s condition factor: weight/length³) between dietary treatment groups (AL and CR) and age control groups (CO and CY) on the x-axis. The y-axis represents Fulton’s condition factor. A Tukey HSD test examining interactions revealed a significant difference between AL and CY (adj. p < 0.0001) and AL and CR (adj. p < 0.001), but not between AL and CO. The difference between CR and CY was highly significant (adj. p < 0.007), whereas the difference between CR and CO was slightly less pronounced (adj. p < 0.03). For interactions that include sex, please refer to Supplementary Table S6, which contains the full statistical details.

**Figure G:** This plot compares relative ovary weight (y-axis, expressed as a percentage of total body weight) across dietary treatment groups (AL, CR) and age control groups (CO, CY) on the x-axis. A Tukey HSD test found no significant differences between most groups, except for a marginal difference between CO and CY females (adj. p = 0.016), and between CR and CY females (adj. p = 0.023).

**Figure H:** Example images of an AL male seahorse (left) and a CR male seahorse (right), illustrating differences in pouch size. The red line represents the diagonal measurement of pouch length. Measurements were performed using ImageJ, with the centimetre paper in the background serving as a reference for scale. Additionally, manual measurements were taken to ensure accuracy and provide a proper comparison.

**Figure I and G:** These figures compare the morphological measurements of seahorses euthanized before the end of the experiment to those that completed the 5-month dietary intervention. This is to show that there were no physiological disadvantages for the seahorses that did not make it to the end of the study. For both length (Figure I) and weight (Figure J) at Day 1, no significant differences were observed between the two groups: length (cm) on Day 1 (ANOVA, non-survivors vs. survivors, p = 0.28) and weight (g) on Day 1 (Kruskal-Wallis, χ² = 0.9, p = 0.33).

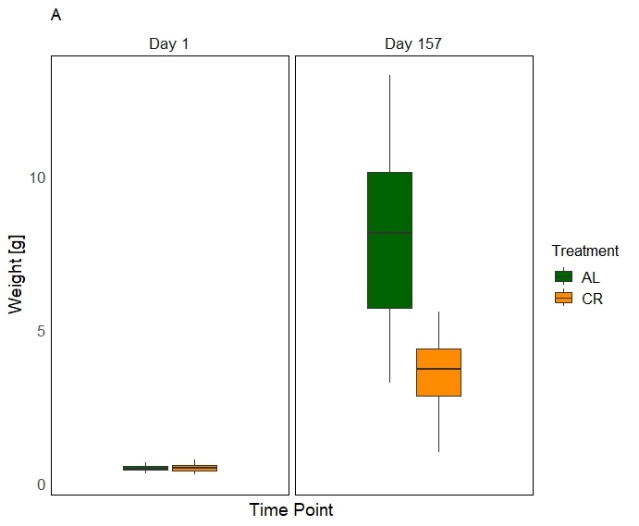

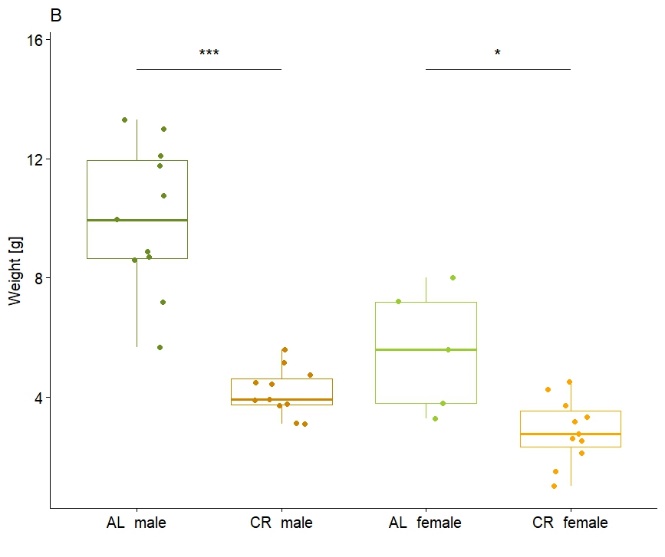

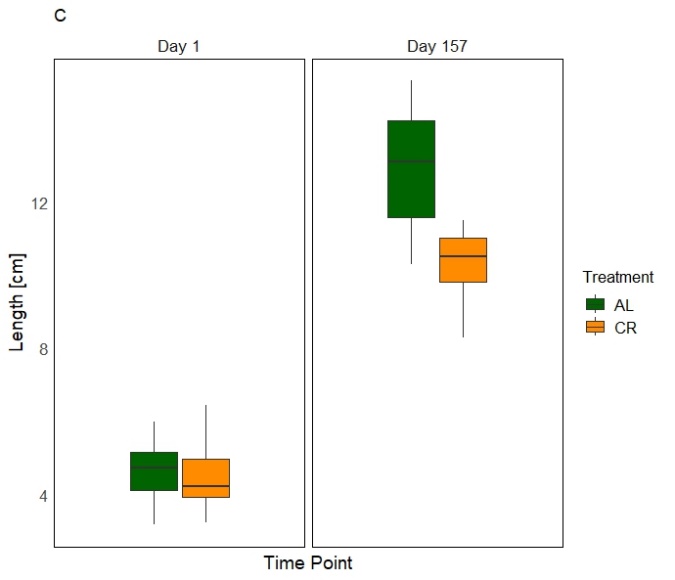

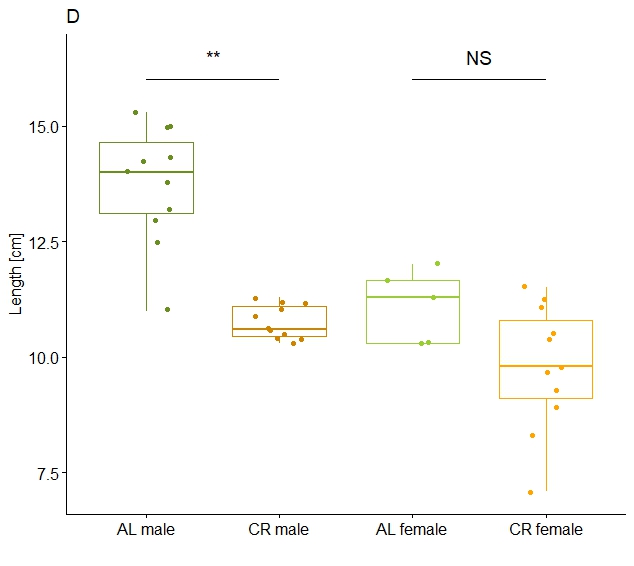

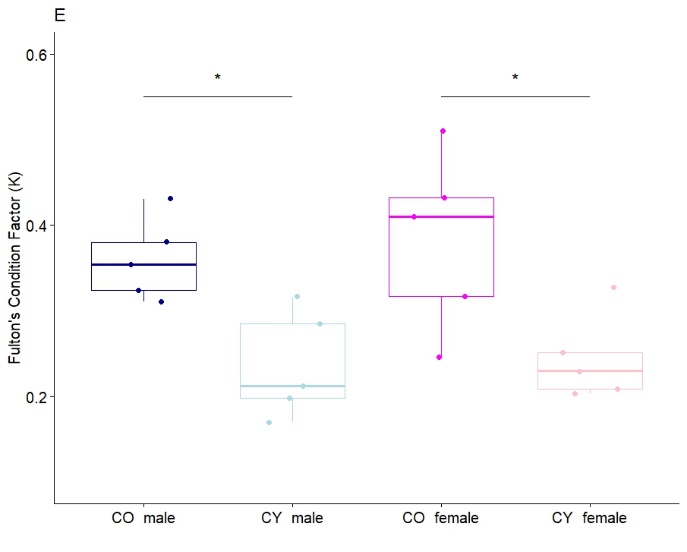

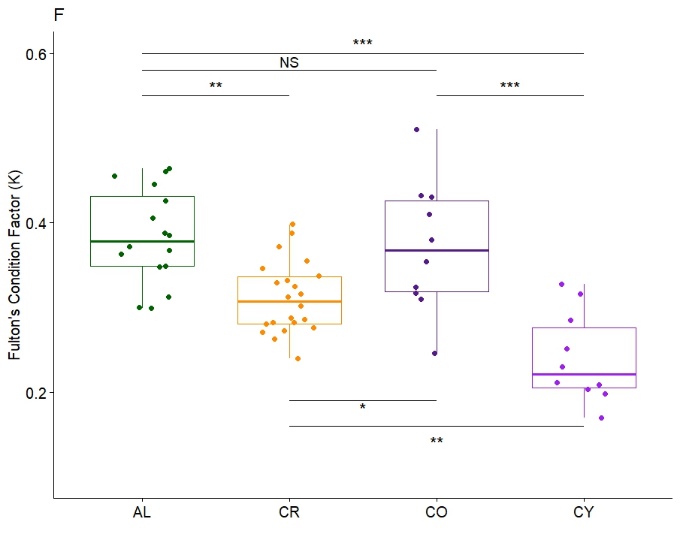

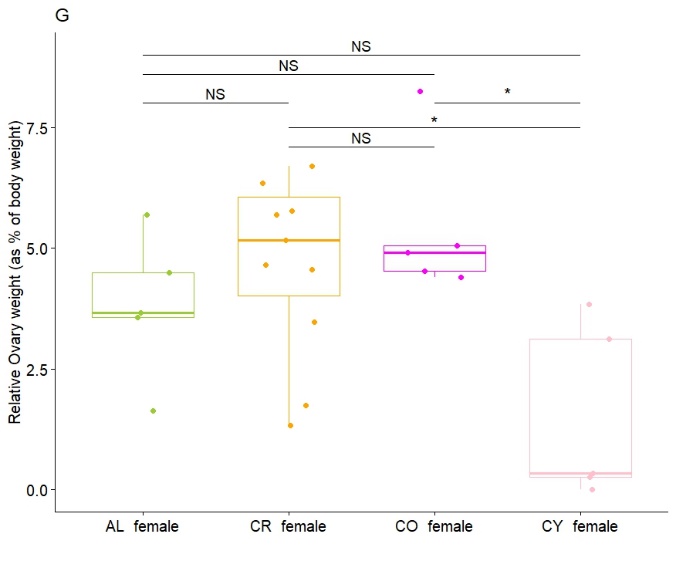

H

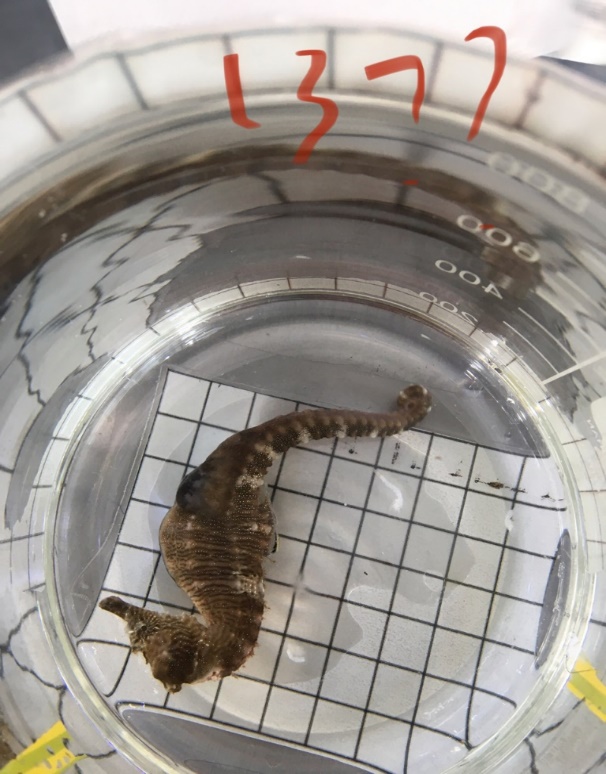

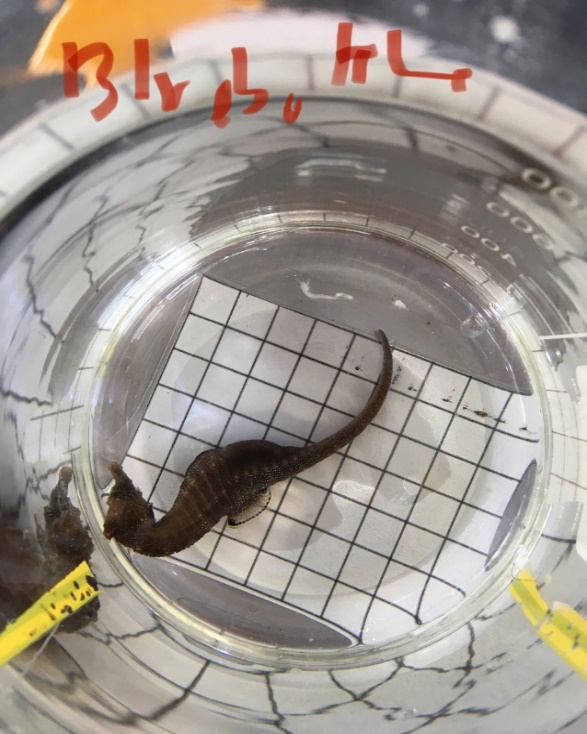

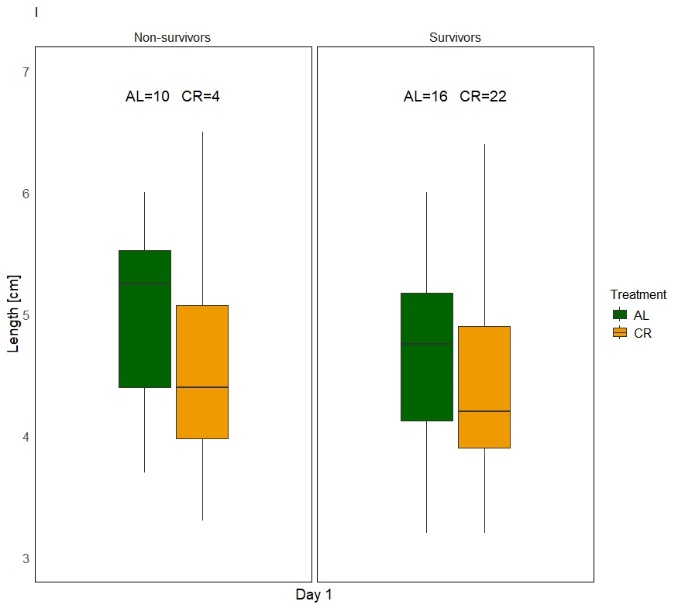

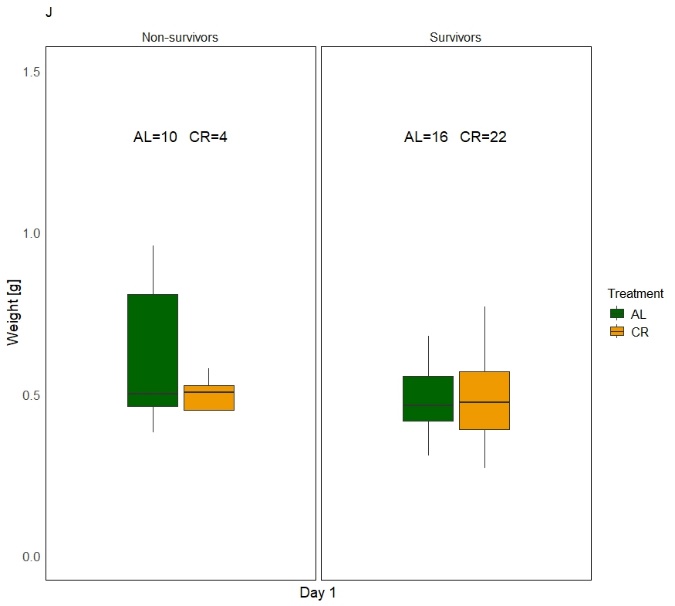

**3.2 Effects of calorie restriction on tissue composition**

**Figure S2:** **Principal Component Analysis (PCA) plot illustrating the separation of organs and treatment groups based on the first two principal components.**

The plot shows the separation of organs across PC1 (x-axis) and PC2 (y-axis). Organs are represented by different symbols: liver (square), head kidney (circle), ovaries (downward triangle), and testes (upward triangle). Treatment groups are color-coded as follows: CR (orange), AL (green), juvenile control (light blue), and adult control (dark blue). PC1 explains 34% of the total variance, with a clear separation of the liver from the other organs.

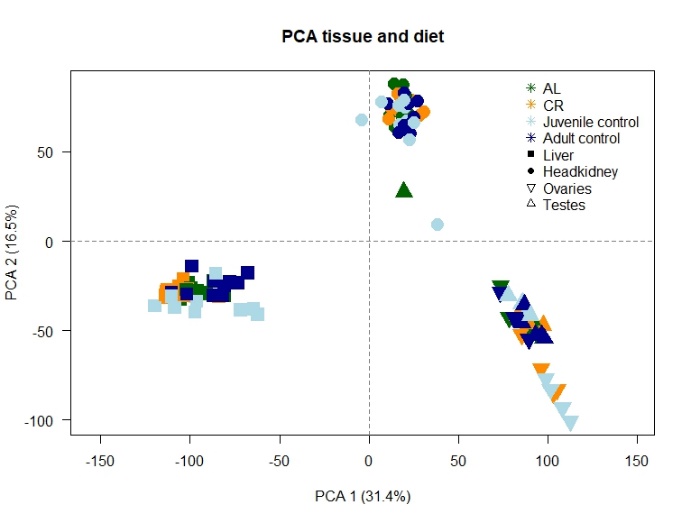

**Table S1:** Quantitative results from the differential gene expression (DGE) analysis (using *limma:voom)* comparing different groups. Only differentially expressed genes below an adj.p.val of 0.05 are showed in the table (one-way analysis of deviance).

| **Contrast comparison** | **Tissue** | **DEGs (adj.p.val < 0.05)** |
| --- | --- | --- |
| AL vs CR | Liver | 940 |
| AL vs CR (female) | Liver | 375 |
| AL vs CR (male) | Liver | 83 |
| AL male vs AL female | Liver | 501 |
| CR male vs CR female | Liver | 83 |
| AL vs CO | Liver | 3195 |
| AL vs CY | Liver | 5408 |
| CR vs CO | Liver | 5003 |
| CR vs CY | Liver | 4292 |
| CO male vs CO female | Liver | 821 |
| CY male vs CY female | Liver | 981 |
| AL vs CO (male) | Liver | 933 |
| AL vs CY (male) | Liver | 4241 |
| CR vs CO (male) | Liver | 2454 |
| CR vs CY (male) | Liver | 3048 |
| AL vs CO (female) | Liver | 1601 |
| AL vs CY (female) | Liver | 1891 |
| CR vs CO (female) | Liver | 2442 |
| CR vs CY (female) | Liver | 601 |
| AL vs CR | Head kidney | 67 |
| AL vs CR (female) | Head kidney | 0 |
| AL vs CR (male) | Head kidney | 4 |
| AL male vs AL female | Head kidney | 973 |
| CR male vs CR female | Head kidney | 1728 |
| AL vs CO | Head kidney | 1566 |
| AL vs CY | Head kidney | 2981 |
| CR vs CO | Head kidney | 2392 |
| CR vs CY | Head kidney | 4106 |
| CO male vs CO female | Head kidney | 153 |
| CY male vs CY female | Head kidney | 66 |
| AL vs CO (male) | Head kidney | 11 |
| AL vs CY (male) | Head kidney | 1674 |
| CR vs CO (male) | Head kidney | 801 |
| CR vs CY (male) | Head kidney | 2846 |
| AL vs CO (female) | Head kidney | 502 |
| AL vs CY (female) | Head kidney | 632 |
| CR vs CO (female) | Head kidney | 558 |
| CR vs CY (female) | Head kidney | 1054 |
| AL vs CR | Testes | 0 |
| AL vs CO | Testes | 0 |
| AL vs CY | Testes | 646 |
| CR vs CO | Testes | 0 |
| CR vs CY | Testes | 345 |
| AL vs CR | Ovaries | 366 |
| AL vs CO | Ovaries | 0 |
| AL vs CY | Ovaries | 4966 |
| CR vs CO | Ovaries | 218 |
| CR vs CY | Ovaries | 0 |

**3.3 Gene Enrichment Analysis Reveals Sex-Specific Fasting Effects on Liver Metabolism**

**Figure S3: Bar plot illustrating sex-biased gene expression in seahorses under ad libitum (AL) and caloric restriction (CR) dietary treatments.**

Genes upregulated in males are represented in blue, while genes upregulated in females are shown in red. The x-axis represents the number of differentially expressed genes with a significant adjusted p-value (< 0.05), corrected using the Benjamini-Hochberg (BH) method. The y-axis displays different organs (liver, head kidney, and gonads, with testes for males and ovaries for females), categorized by CR or AL treatment groups.

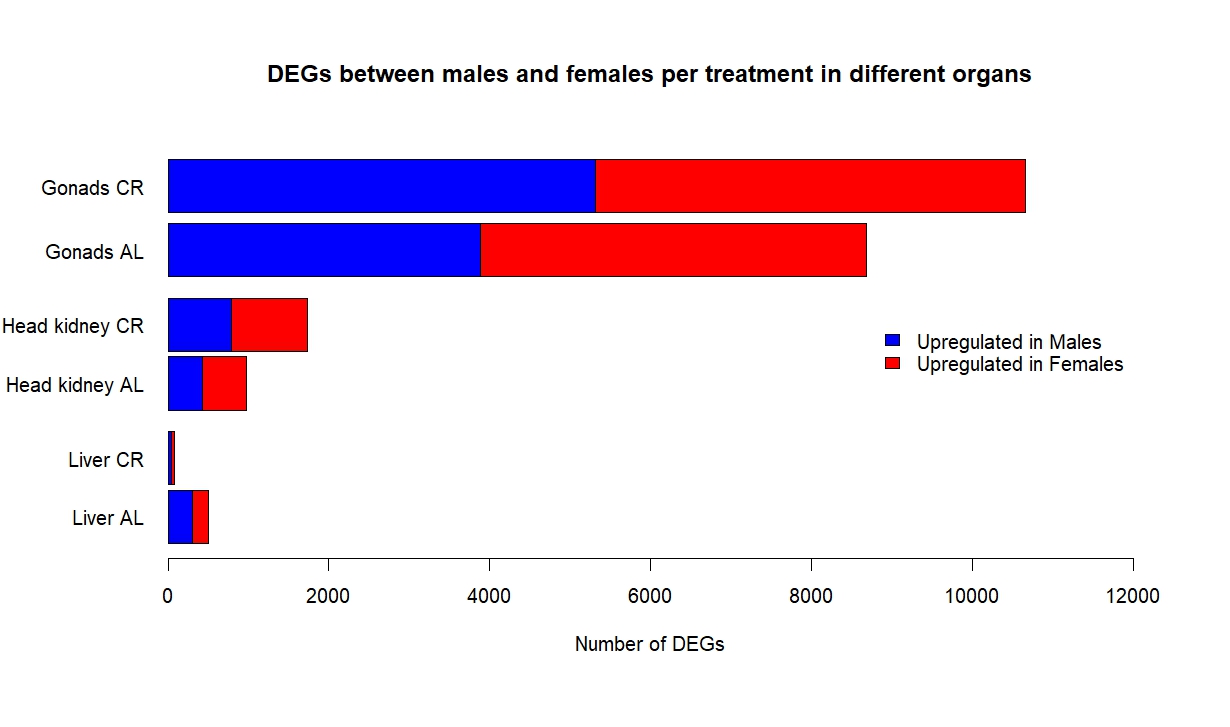

**3.5 Calorie restriction promotes greater gut microbial diversity**

**Figure S4: Microbial diversity and composition across treatment and age groups in seahorses.
A)** Boxplots of alpha diversity (Shannon index) for treatment groups (ad libitum [AL] and caloric restriction [CR]) and age groups (control old [CO] and control young [CY]). The x-axis represents the groups, while the y-axis shows Shannon richness. No significant differences were observed between diet or age groups (see Table S6).

**B)** Bar plot showing the relative abundance of microbial taxa at the phylum level, including the most abundant genus, Vibrio. Panels are categorized by diet (AL and CR) and age (CO and CY), with further division by sex (male and female seahorses) within each subcategory.

**C)** Non-metric multidimensional scaling (NMDS) plot based on the Bray-Curtis dissimilarity matrix, comparing microbial communities across all group variables: diet (AL in green and CR in orange) and age (CO in dark blue and CY in light blue).

**D)** NMDS beta diversity plot based on Bray-Curtis dissimilarity, comparing dietary treatment groups (AL and CR) by sex. Each dot represents an individual, and 95% confidence ellipses indicate group variability, with ellipses based on standard error. Arrows highlight microbial taxa significantly associated with community structure (p < 0.001), pinpointing key indicator species for each group.

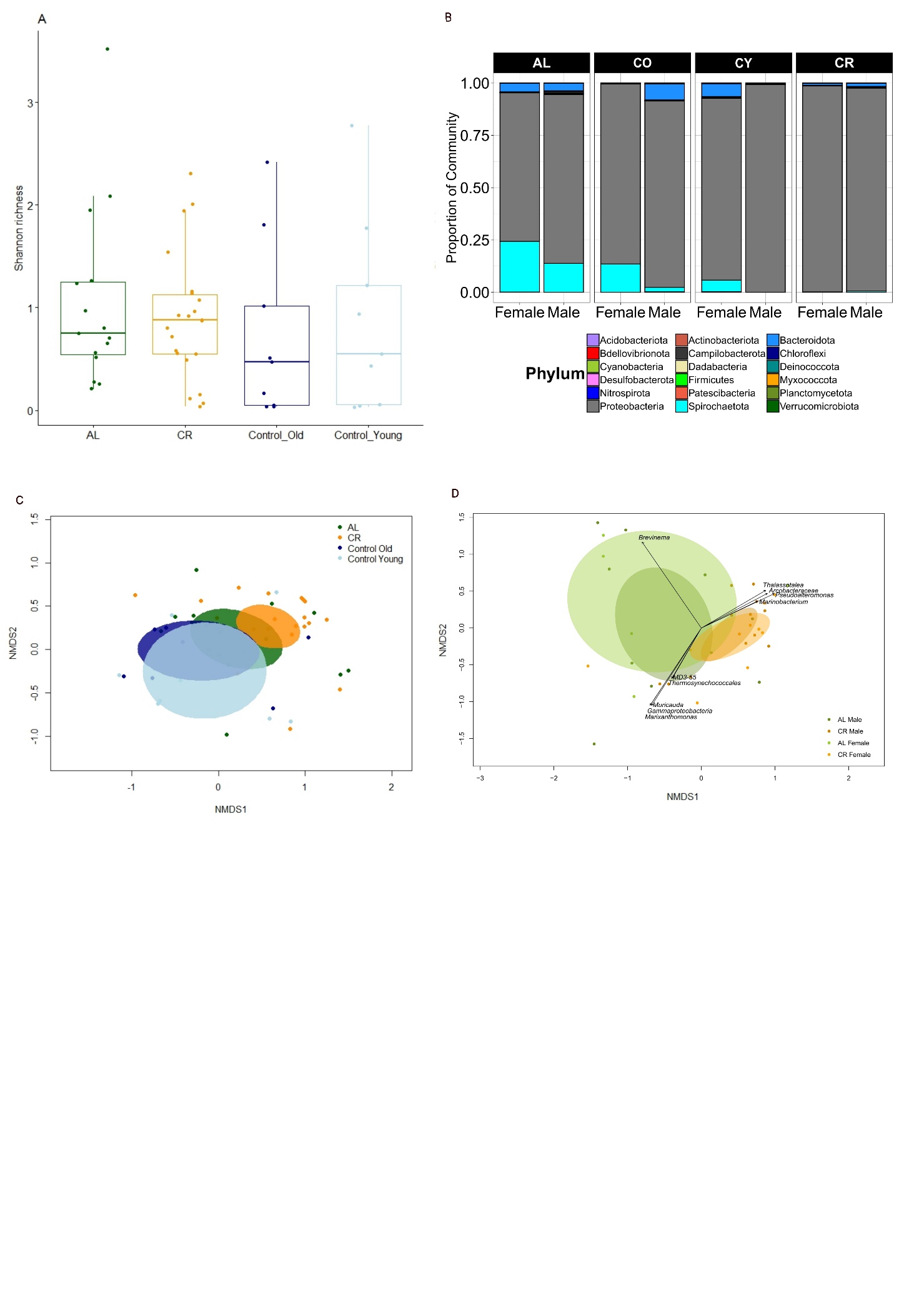

**3.6 Correlation analysis between gut microbiota and transcriptomic data**

To assess potential associations between microbiota composition and host transcriptomic profiles, we performed a Mantel test comparing the 205 bacterial genera identified in our 16S rRNA sequencing data with the normalized transcriptomic data from liver, head kidney, testes, and ovaries (Table S6). The Mantel test, which evaluates correlations between two distance matrices using the Bray-Curtis method, revealed no significant associations for the liver and head kidney. However, significant correlations were detected for the ovaries (p = 0.01) and testes (p = 0.013), suggesting a potential relationship between microbial composition and transcriptional variation in these reproductive tissues.

To further explore these associations, we conducted a Multidimensional Scaling (MDS) analysis. In the ovaries, MDS analysis of the AL and CR treatment groups identified seven significant bacterial taxa (p < 0.05), all of which were associated with the AL diet: *Altererythrobacter, Flavobacterium, Portibacter, Spongiimicrobium, Pontivivens, Paraspirulinaceae,* and *Jannaschia*. Additionally, MDS analysis between microbiota and transcriptomic data identified 781 significant associations (p < 0.01) in the ovaries. Gene Ontology (GO) and KEGG pathway enrichment analysis of these associated genes revealed functional categories related to catalytic activity (GO:0003824), lysosome function (KEGG:04142), biosynthesis of unsaturated fatty acids (KEGG:01040), cytoplasm (GO:0005737), and membrane-associated functions (GO:0016020), amongst other pathways.

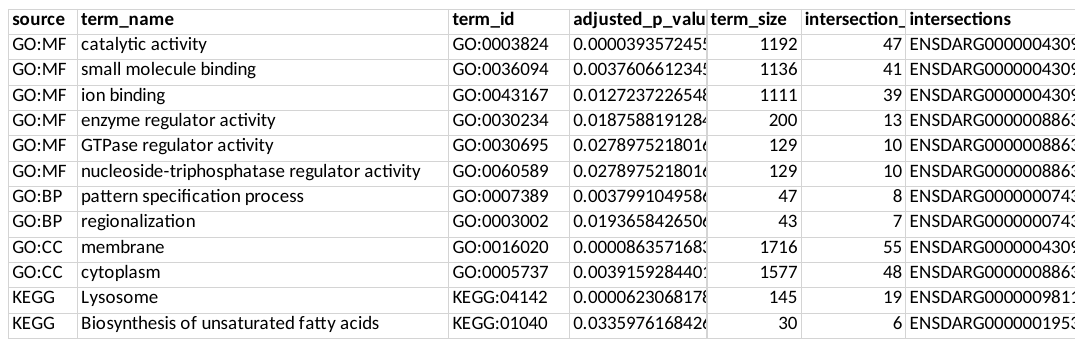

In the testes, we identified five significant bacterial taxa (p < 0.05): *Vibrio, Thalassotalea, Brevinema, Pseudoteredinibacter,* and *Colwellia*. MDS visualization indicated that *Thalassotalea, Pseudoteredinibacter*, and *Colwellia* were positioned in the direction of the CR treatment group (Figure S5), and *Brevinema* for AL, mirroring results in Figure 4B. Gene enrichment analysis of the 345 significant genes (p < 0.01) identified in the testes MDS analysis revealed a single overrepresented Gene Ontology (GO) term: membrane (GO:0016020).

Lastly, we specifically examined immune-related transcriptomic variation and its correlation with microbial taxa in both tissues. In the ovaries, no significant relationships were detected (Table S6). However, in the testes, we identified ~20 immune-related genes that exhibited significant correlations with microbiota composition (p < 0.05), as visualized in Figure S5. While immune-related genes did not cluster strongly within the same region, FADD, hhla2a, and tcima showed some alignment with *Pseudoteredinibacter, Thalassotalea* and *Colwellia* for CR. In contrast, *Brevinema* was positioned toward the AL treatment group, where a greater number of immune-related genes were located, including irf10, mhc1zca, cd74, lck, FRK, and others (Figure S5). *Vibrio* was gravitated to an intermediate space between AL and CR, with only marchf8 appearing in the same region. These spatial patterns suggest potential associations between specific bacterial taxa and immune-related transcriptional variation, particularly in AL-treated samples, where *Brevinema* co-occurs with multiple immune genes. These findings suggest that interactions between microbial communities and host immune pathways may differ between reproductive tissues and dietary treatments, with potential implications for immune-microbiota crosstalk.

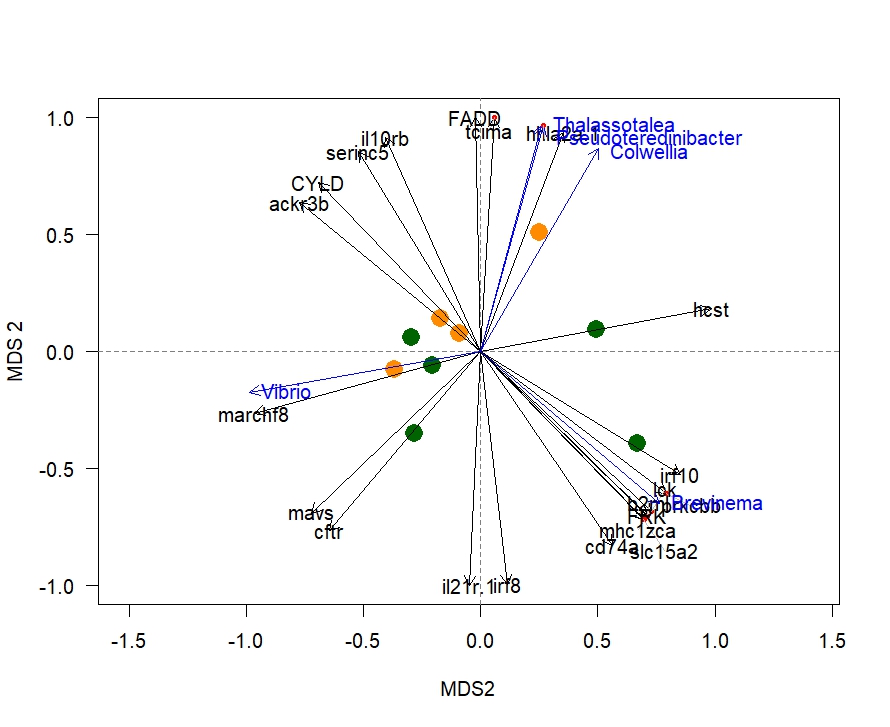

**Figure S5: Multidimensional Scaling (MDS) analysis of microbial composition and gene expression in male Hippocampus erectus testes tissue across dietary treatments.**

MDS is an ordination method that represents high-dimensional data in a reduced space, preserving the relative distances between samples to visualize similarities and differences. Here, individual samples from the CR group are shown in orange, while those from the AL group are shown in green. Black arrows indicate significant immune-related genes, while blue arrows represent significant bacterial taxa. These vectors were determined using *envfit*, which fits environmental variables (in this case, gene expression and microbial taxa) onto the MDS ordination space to identify significant associations. The arrows represent variables that were significantly correlated with the ordination space (p < 0.05), as calculated using the function *envfit(micronMDS, t(OTU_exp_testes))*. Only significant vectors *(xfit$vectors$pvals < 0.05)* are displayed. The direction of the arrows suggests the gradient along which specific genes and taxa are associated with sample distributions.
